# De novo identification of protein domains in cryo-electron tomography maps from AlphaFold2 models

**DOI:** 10.1101/2024.11.21.623534

**Authors:** Yi Lu, Guanglin Chen, Fei Sun, Yun Zhu, Zhiyong Zhang

## Abstract

In situ cryo-electron tomography (cryo-ET) enables visualization of protein complexes within their native cellular environments. However, sub-tomogram averaging (STA) often fails to achieve resolutions sufficient for atomic model building based on side-chain information, hindering accurate protein identification and assignment. To address this challenge, we developed DomainSeeker, a computational workflow that integrates AlphaFold2-predicted structures with experimental data to identify protein domains in cryo-ET density maps. DomainSeeker partitions AlphaFold2-predicted structures into domains using clique analysis based on predicted aligned errors and then fits these domains into target densities segmented from input cryo-ET maps. Each domain-density fit is evaluated globally and locally to derive prior probabilities, which are subsequently integrated with complementary data, such as cross-linking mass spectrometry (XL-MS), through Bayesian inference to compute posterior probabilities. Applying DomainSeeker’s prior probability analysis to densities from mouse sperm microtubule doublets revealed substantially higher recognition accuracy and resolution robustness compared with existing approaches. Incorporating XL-MS data for yeast nuclear pore complex densities further enhanced both recognition performance and confidence. Finally, we implemented an interactive ChimeraX plugin to facilitate user access. Together, DomainSeeker overcomes a key bottleneck in in situ structural biology, providing a robust and accessible tool for visual proteomics to uncover the structural and functional organization of native cellular assemblies.

## Introduction

Over the past decades, cryo-electron microscopy (cryo-EM) has enabled the visualization of biomolecular complexes at near-atomic resolution^1^, while cryo-electron tomography (cryo-ET) has extended this capability to determine protein structures within their native cellular environments^2,3,4^. Recent cryo-ET studies have provided critical in situ insights into diverse macromolecular assemblies, offering understanding of their functional mechanisms^5,6,7^. By revealing the organization and interactions of the intracellular proteome, in situ cryo-ET has driven the emergence of visual proteomics^8,9^, linking molecular structure to cellular architecture.

However, in situ structures resolved by sub-tomogram averaging (STA) often fail to reach the resolution required for atomic model building^10,11^. Consequently, for regions with invisible sidechains of in situ cryo-ET maps, identifying proteins and constructing atomic models directly from the density maps alone remain infeasible^12^. For these densities, candidate proteins can instead be inferred from mass spectrometry (MS) data^13^, whose experimentally determined or computationally predicted structures are fitted into the target densities to identify the best match. The key challenge lies in obtaining reliable structural models for a large number of proteins. Recent advances in protein structure prediction have made this possible^14,15,16,17,18^, particularly AlphaFold2, which achieves near-experimental accuracy in a majority of cases^15^. The AlphaFold Protein Structure Database (AFDB) now provides over 200 million predicted models across numerous species^19^, allowing rapid and automated retrieval of candidate structures for density fitting.

Several studies have adopted this strategy. Fitting AlphaFold2-predicted models using Colores^20^ identified multiple novel microtubule inner proteins (MIPs) within mouse sperm axonemal microtubules^21^. Another method, DomainFit, demonstrated domain identification in cryo-EM density maps at 6–8 Å resolution^22^. However, existing approaches remain limited by low efficiency, complex implementation, and suboptimal accuracy. Moreover, no established framework yet integrates complementary experimental evidence—such as cross-linking mass spectrometry (XL-MS)^23^—into the identification process. These gaps underscore the need for a robust, accurate, and versatile workflow for protein identification in situ.

To address this, we developed DomainSeeker, an efficient computational workflow and software tool for identifying protein domains in cryo-ET density maps by integrating AlphaFold2-predicted structures with complementary experimental data (e.g., XL-MS). DomainSeeker comprises four modules: (1) partitioning AlphaFold2 models into domains via clique analysis of predicted aligned errors (PAE)^24,25,26^;(2) fitting these domains to segmented target densities in ChimeraX^27,28^; (3) globally and locally evaluating the quality of each domain–density fit; and (4) calculating posterior probability via Bayesian inference to integrate all evidence for enhanced recognition capability and reliability.

To validate prior probabilities, we segmented 26 densities from a sperm microtubule doublet cryo-ET map (~6.5 Å)^29^ based on atomic models. DomainSeeker successfully recognized all corresponding domains, maintained robustness at 10 Å after low-pass filtering, identified a previously unmodeled protein, and correctly detected 17 out of 20 auto-segmented densities—outperforming existing methods. For posterior validation, DomainSeeker analyzed 31 yeast nuclear pore complex (NPC) densities^30^, correctly identifying 18 regions based on prior probabilities alone and 4 additional regions after incorporating XL-MS data. We also developed a graphical ChimeraX plugin to streamline computational workflows and result inspection.

Together, DomainSeeker provides a powerful, user-friendly framework for cryo-ET–based visual proteomics, bridging predicted protein structures and experimental density maps to reveal new structural and functional insights in native cellular environments.

## Results

### Workflow of DomainSeeker

DomainSeeker is a computational framework designed to identify protein domains within cryo-ET density maps by integrating AlphaFold2-predicted structures with complementary experimental data such as XL-MS^23^. As illustrated in Figure 1, the workflow comprises four modules.

**Figure 1.**
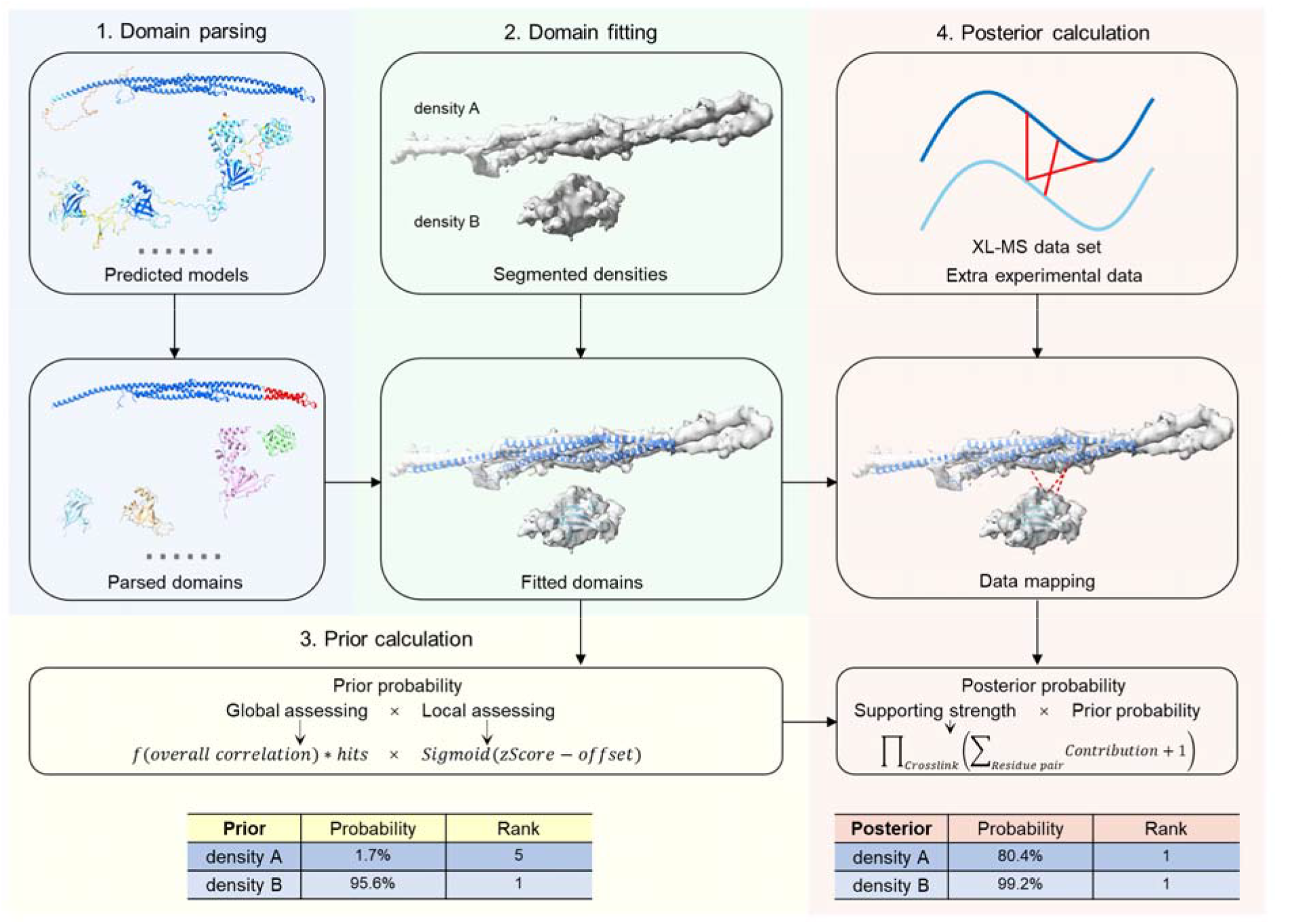
Workflow of DomainSeeker. Module 1: partition predicted models into domains. Module 2: fit all domains into every segmented density. Module 3: calculate prior probabilities using both global and local fitting assessments. Module 4: calculate posterior probabilities using Bayesian inference.

Module 1: Domain parsing. Although AlphaFold2 reliably models compact domains, it can incorrectly predict inter-domain orientations and positions^31^. Candidate proteins are typically obtained from mass spectrometry (MS) data. When MS data are unavailable, candidates can be selected from the complete proteome of the target organism. DomainSeeker can automatically retrieve predicted structures and corresponding PAE files from the AFDB and partition each protein into domains through PAE-based clique analysis.

Module 2: Domain fitting. The input cryo-ET map is first segmented into target density regions that potentially correspond to individual proteins or domains, either manually or automatically in ChimeraX^27^. Segments with poor signal-to-noise ratios are excluded to ensure data quality. For each target density, ChimeraX is then used to identify candidate fitting positions for every domain, with each position representing a local maximum in the cross-correlation between the domain model and the target density.

Module 3: Prior calculation. To quantify confidence in each domain placement, DomainSeeker computes two complementary metrics: global and local fitting estimations. The global estimation calculates the probability that a fitting position is correct among all potential matches for the domain within the target density based on overall structural correlation. The local estimation evaluates the correlation between overlapping regions of the domain and the density. These two measures are integrated to yield a prior probability, reflecting the confidence that a specific domain–density fit is correct.

Module 4: Posterior calculation. Posterior probabilities are computed by integrating all available evidence through Bayesian inference. Currently, DomainSeeker has incorporated XL-MS data by mapping cross-links onto candidate structural states and evaluating compliance with distance constraints. The supporting strength of the input data for each state is calculated by summing contributions across positions for the same cross-link and multiplying contributions from different cross-links, thereby weighting fits according to experimental consistency.

Finally, a ChimeraX plugin was developed for DomainSeeker to facilitate both the computational processes and the following result inspection. This helps users to efficiently inspect results and assemble initial structural models with confidence.

In this work, we not only established a complete pipeline to process cryo-ET reconstructed maps but also developed several key algorithms to execute specific tasks within the established pipeline—including domain parsing based on PAE clique analysis, candidate fitting position screening, and prior/posterior probability calculation.

### Key algorithms

#### Parsing domains based on PAE-derived cliques

To partition AlphaFold2-predicted structures into domains, DomainSeeker uses the PAE, which reflects interdomain flexibility, together with the per-residue local distance difference test (pLDDT), which reports local confidence^19^. Using the protein with UniProt ID Q9D485 as an example (Fig. 2a), we constructed a residue graph in which nodes (residues with pLDDT ≥70) are connected if their average PAE is <5. Fully connected subgraphs, or cliques, represent rigid structural entities. Small cliques (<5 residues), typically corresponding to flexible loops, are excluded. Connected cliques (sharing ≥50% residues or with >60% PAE-linked edges) are merged into clique groups, which together define structural domains. Groups with fewer than 40 residues (Fig. 2a, gray nodes in the clique graph) are removed, as they often fit nonspecifically into many densities or correspond to short, flexible regions that reduce fitting reliability.

**Figure 2.**
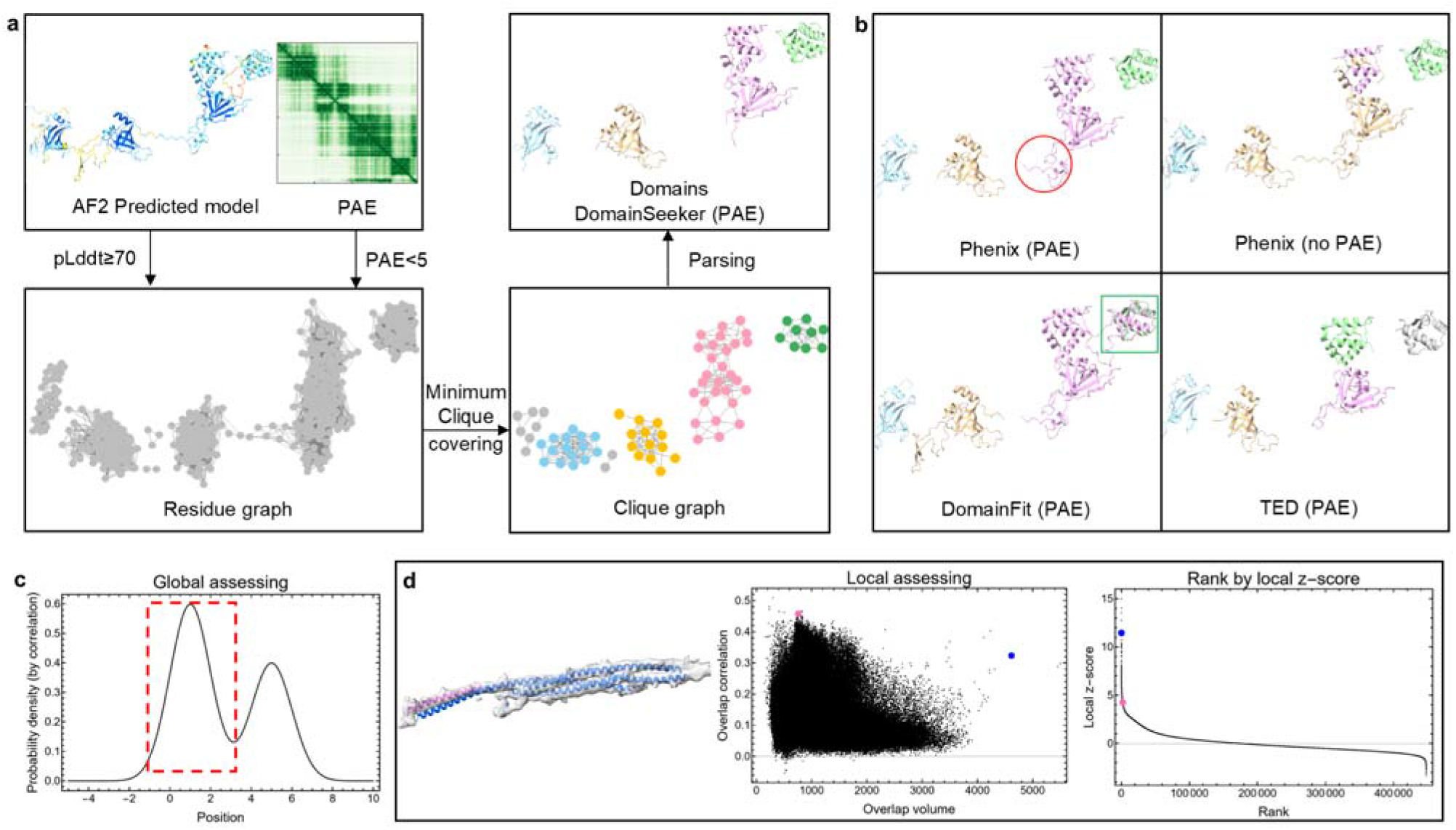
Key algorithms. (a) Domain parsing workflow in DomainSeeker (sample protein Q9D485). Residues with pLDDT <70 or mutual PAE >5 are excluded to construct a residue graph. Minimum clique covering is applied, with the resulting cliques as nodes in a new clique graph. Cliques are linked if they share >50% residues or have >60% PAE edges; interconnected cliques form domains. (b) Domains partitioned by Phenix, DomainFit and TED are color-coded. The red circle: a flexible loop in Phenix but absent in DomainSeeker. The green box: a region incorporated into the pink domain by DomainFit. The gray region in the TED result was not assigned as a domain due to insufficient consensus among the three methods (Chainsaw, Merizo, and UniDoc). (c) The probability density curve of fitting positions (correlation-derived), with a red dashed box highlighting a local maximum peak. (d) Left: domain corresponding to density A (blue) and a short helix (pink) fitted to density A. Middle: overlap region volume vs. correlation (blue: the domain corresponding to density A; pink: the short helix). Right: Rankings vs. local z-scores (the blue domain ranks 5th).

This strategy enables DomainSeeker to robustly decompose predicted structures into domains across diverse architectures and PAE patterns. As shown in Fig. 2a, the protein is partitioned into four well-defined domains consistent with the PAE plot: each domain shows high internal reliability, while interdomain orientations remain uncertain. In comparison (Fig. 2b), Phenix offers two domain parsing methods with its phenix.process_predicted_model utility^32,33^. Its PAE-based method yields four domains matching the expected count but still includes a flexible terminal loop (the red circle, Fig. 2b). The default method, which partitions structures solely based on 3D geometry, performs worse: it incorrectly merges the two middle domains and splits the third, impairing downstream fitting and scoring. DomainFit^22^, which reads Phenix outputs, merges the third and fourth domains (the green box, Fig. 2b), potentially compromising identification when their relative orientations are inaccurate.

AFDB also provides domain annotations from TED^34^, which identifies domains by consensus of three deep learning methods—Chainsaw^35^, Merizo^36^, and UniDoc^37^. On the same protein (Fig. 2b), TED split DomainSeeker’s third domain into two smaller subdomains and left a gray region unassigned due to insufficient consensus. Although this division is functionally justified, DomainSeeker pursues a different objective: it defines domains as rigid structural units that retain as many distinctive features as possible, while ensuring structural accuracy and rigidity. Its exclusion of flexible termini through clique-based filtering could further improve fitting precision.

#### Considering all fitting positions of each domain in each density

DomainSeeker identifies all plausible placements of each domain within every segmented density using ChimeraX. The algorithm initializes each domain at random positions within the target density and performs gradient ascent optimization to locate local cross-correlation maxima. These maxima are clustered by spatial proximity, with ChimeraX reporting both the cross-correlation at each cluster centroid and the number of contributing maxima (“hits”).

Because the correct placement is not always the top-ranked solution, methods that consider only the highest-scoring fits may miss the correct domains. DomainSeeker mitigates this by recording all fitting positions for each domain–density pair and subsequently evaluating their prior probabilities, reducing the likelihood of omissions due to incorrect fit selection.

#### Evaluating prior probabilities using global and local fitting assessments

For each candidate placement, DomainSeeker quantifies correctness probability using two complementary metrics: global and local fitting assessments.

The global metric estimates the likelihood that a placement is correct among all candidates of this domain within the target density. As illustrated in Fig. 2c, local maxima of the cross-correlation correspond to probability peaks in six-dimensional translational–rotational space. The probability of the correct placement within each peak (red dashed region) is approximated as the product of the peak’s maximum probability density and its hypervolume, which scales with the number of convergent hits. Thus, global confidence increases with both cross-correlation strength and hit count.

While global correlation captures overall shape similarity, it may underestimate correct placements when the target density represents only part of a domain or, conversely, when the density extends beyond the domain. To assess local structural compatibility, we apply a Laplacian filter^38^ to enhance intra-domain contrast and compute cross-correlation only within overlapping regions of the domain and density. However, overlap correlation alone may favor small fragments that fit broadly across densities. For example (Fig. 2d), a small helix (pink) shows high overlap correlation yet is less representative than the larger domain (blue). We found that by plotting fitted domains against overlap volume and correlation, well-fitted domains stand out over others with similar overlap volumes. To quantify this observation, DomainSeeker computes a local z-score, defined as the vertical deviation (in standard deviations) of each point from the local mean curve of correlation over volume. The blue domain in Fig. 2d exhibits a markedly higher local z-score, consistent with the correct placement.

Global and local metrics are integrated to yield the prior probability, representing the likelihood that a given domain–density fit is correct. Other tools that rely solely on cross-correlation perform only this global assessment.

#### Evaluating posterior probabilities based on Bayes’ Theorem

DomainSeeker refines prior probabilities by integrating additional experimental evidence through Bayesian inference. Currently, XL-MS data are supported. For each structural state, the posterior probability is proportional to its prior probability multiplied by a scaling factor derived from the XL-MS dataset. This factor is calculated as the product of contributions from individual cross-links, where each contribution equals the summed compliance across all residue pairs satisfying distance constraints.

Detailed derivations are provided in the Methods. To our knowledge, DomainSeeker is so far the only tool for protein domain identification in cryo-ET density maps that can directly incorporate experimental data into probabilistic inference.

The impact of XL-MS integration depends in part on data quality. DomainSeeker does not explicitly model the false discovery rate (FDR)^39^; instead, its design inherently limits the influence of false-positive cross-links, that is, a cross-link can only affect the posterior probability of a candidate if it connects to a high-confidence partner. This is unlikely for false positives in datasets with reasonably low FDR. That said, MS data are intrinsically noisy, and distinguishing every false-positive cross-link from a true one remains impractical without additional information.

### Identifying domains using prior probabilities

To validate the prior probabilities estimated by DomainSeeker, we generated a test set using a cryo-ET map of the mouse sperm microtubule doublet (EMD-35230, ~6.5 Å resolution) and its corresponding atomic model (PDB 8I7R)^29^. Among the 36 MIPs, 16 either lack well-defined structured domains or consist primarily of extended single helices, which require alternative identification strategies. We therefore selected 26 domains from the remaining 20 MIPs for analysis. A total of 4,303 proteins from the MS dataset were used as candidate inputs.

For comparison, we applied DomainFit to the same dataset. DomainFit uses phenix.process_predicted_model for domain parsing, calculates the top-fitting position for each domain in each density using ChimeraX, and ranks candidates by p-values of the corresponding correlations, reflecting how distinct the top fit is among all candidates. Domains parsed by DomainSeeker and Phenix are shown in Figure S1. Overall, DomainSeeker’s partitioning facilitates more accurate domain identification.

#### Performance on model-based densities

We first evaluated DomainSeeker’s performance using densities segmented based on reference domains. To segment the 6.5 Å density map, we simultaneously loaded the map and corresponding domains into ChimeraX, colored regions within 5 Å of the 26 target domains using the Color Zone tool, and extracted them from the original map with the Split Map tool. Test densities at 10 Å resolution were generated with the *molmap* command in ChimeraX.

In all test results presented in this study, a domain placement is considered correct if the assigned domain corresponds to the annotated protein and aligns well with the reference model.

Figure 3a compares the recognition accuracy of DomainSeeker (cyan) and DomainFit (orange), and detailed rankings and probabilities are provided in Figure S2. The left and middle bar groups represent densities at 6.5 Å and 10 Å, respectively.

**Figure 3.**
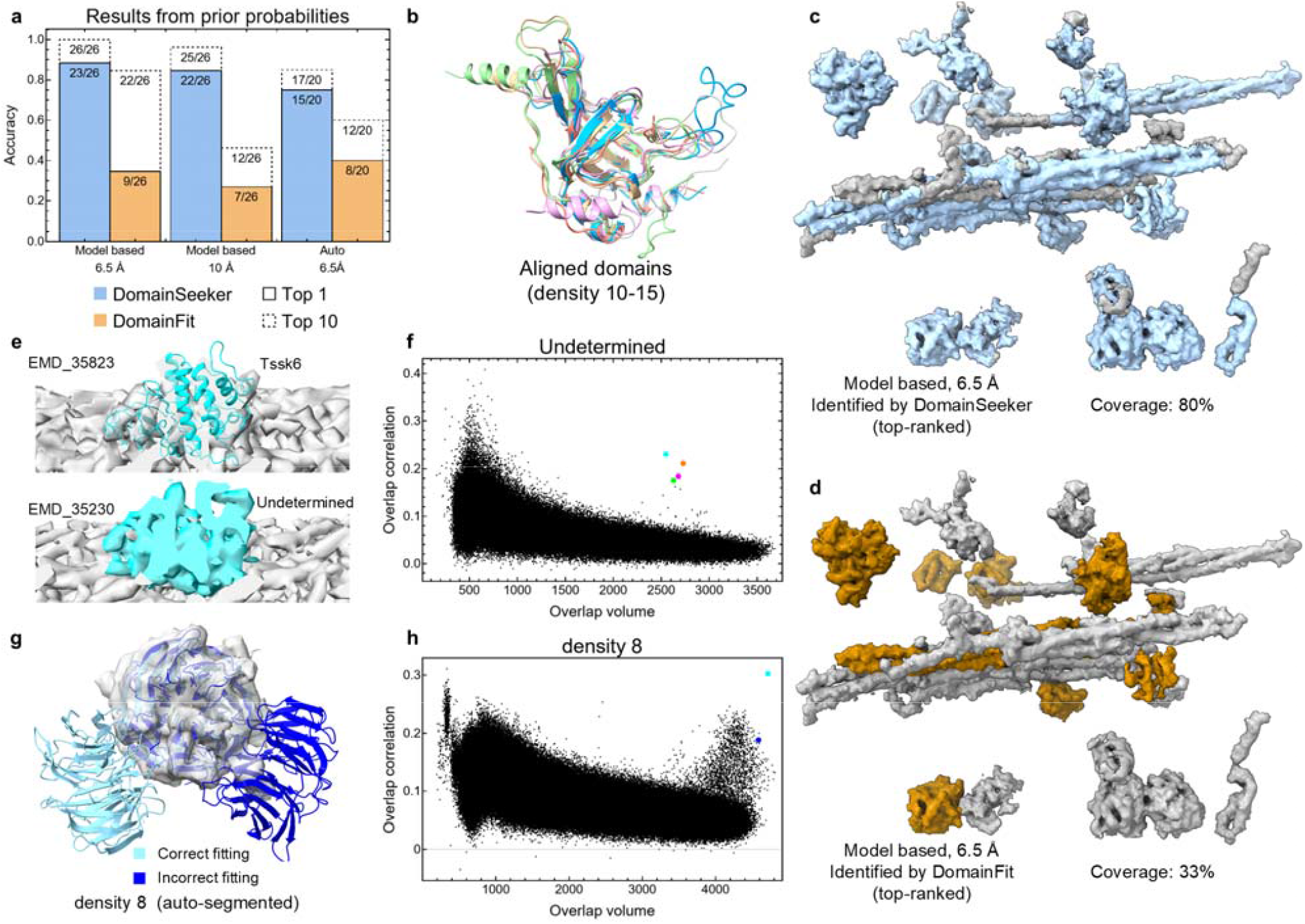
Performance on densities segmented from the mouse sperm microtubule doublet. (a) Recognition accuracy of DomainSeeker (cyan) and DomainFit (orange) on model-based densities (6.5 Å and 10 Å) and auto-segmented densities (6.5 Å). The solid lines represent the proportions of correct domains ranked first, while the dashed lines indicate the proportions of those ranked within top 10. (b) Aligned structures for domains corresponding to densities 10–15. (c) Regions identified by the top-ranked results of DomainSeeker in the model-based densities (6.5 Å). The coverage ratio is 80%. (d) Regions identified by the top-ranked results of DomainFit in the model-based densities (6.5 Å). The coverage ratio is 33%. (e) Tssk6 in cryo-EM/single-particle analysis (SPA) map (EMD_35823) and unmodeled region outside A09 in in situ map (EMD_35230). (f) 2D scoring plot for unmodeled density. Colored points: top four local z-score ranks. Cyan point: Tssk6. (g) The correct fit (cyan) and the incorrect top fit (blue) of the correct domain in density 8 (auto-segmented). (h): Overlap volume vs. correlation of density 8. The correct fit (cyan) and the incorrect top fit (blue).

At 6.5 Å, by combining the global and local scoring (Figure S3) in the prior calculation, DomainSeeker identified the correct domain as the top candidate in 23 of the 26 densities, with the remaining three ranking within the top five (Fig. S2a, Table S1). Of these 26 domains, six structurally analogous domains—derived from homologous proteins Efhc1 and Efhc2—are highlighted in Figure 3b (densities 10-15). For these, DomainSeeker consistently prioritized the correct fit, demonstrating robust discrimination of structurally analogous targets. In contrast, DomainFit ranked the correct domain first in only 9 cases and failed to distinguish the similar domains (densities 10–15). Figures 3c–d illustrate regions identified by the top-ranked results of DomainSeeker and DomainFit in the 6.5 Å dataset, with coverage ratios of 80% and 33%, respectively.

When the densities were low-pass filtered to 10 Å, by integrating the global and local scoring (Figure S4), DomainSeeker maintained nearly identical performance (Fig. S2b, Table S1), whereas DomainFit exhibited substantial degradation. Together, these results demonstrate that DomainSeeker achieves high recognition accuracy and remains robust even at 10 Å resolution.

#### Additional assessment of identification

We carried out additional analyses to assess the contribution of the local and global scoring components, the impact of domain parsing strategies, and the sensitivity to key parameters. To make these extensive evaluations computationally feasible, we first confirmed that a subset of 400 candidate proteins yielded nearly identical rankings as the full set of over 4,000 proteins on the sperm MIP dataset (Fig. S5). All analyses in this section were therefore performed using this reduced set.

Ablation experiments (Fig. S6) showed that the local z-score dominates identification accuracy, with the global correlation providing a complementary yet non-negligible contribution. The ranking performance of TED-derived domains is overall similar to that of DomainSeeker (Fig. S7), although DomainSeeker’s larger rigid domains provided richer structural features, leading to more accurate identification. Systematic sensitivity analysis of domain parsing (Fig. S8), fitting and scoring parameters (Fig. S9) confirmed that identification accuracy is robust to moderate parameter variations. For domain parsing, the current default parameters achieve satisfactory performance. We note that min_domain_size and max_domain_size have a moderate impact on the results (Fig. S8f,g). Users are advised to adjust these two parameters based on the volume of the target density for optimal performance and efficiency.

#### Identification of a new protein

By comparing the in situ cryo-ET map (EMD-35230) with the 3.5 Å cryo-EM/single-particle analysis (SPA) map (EMD-35823)^40^, we identified an unmodeled density adjacent to the outer side of protofilament A09. Figure 3e shows the segments of the mouse sperm microtubule doublet in EMD-35823 (top) and EMD-35230 (bottom). In the cryo-EM/SPA map, this region is annotated as Tssk6, but it was not modeled in the in situ map due to insufficient supporting evidence.

Applying DomainSeeker to this unmodeled density, Tssk6 and three structurally similar proteins were successfully fitted. In the 2D scoring plot (Figure 3f), Tssk6 appears as a cyan point in the upper-right corner, with the top four domains exhibiting comparable local z-scores. With a prior probability of 56.98%, Tssk6 was identified as the top candidate, enabling accurate assignment of the correct structure to this previously unmodeled density.

#### Performance on auto-segmented densities

The previous analysis focused on model-based segmented densities. In practice, unmodeled cryo-ET maps lack reference structures, so automatically segmented densities may not precisely match target domains in size. To assess DomainSeeker’s performance under these conditions, we automatically segmented the 6.5 Å density map using ChimeraX and selected 20 densities that fully or partially covered 20 target domains. Six domains were excluded because they are Tektin-like or elongated, making them challenging for automated segmentation.

After the prior calculation combining global and local results (Fig. S10), DomainSeeker correctly identified 17 of the 20 densities, with 15 ranked first (Fig. 3a, Fig. S2c, and Table S1), significantly outperforming DomainFit. In density 8 (Figure 3g), ChimeraX incorrectly assigned the top fit: the correct domain is shown in cyan, and the incorrect top fit is shown in blue. In the 2D scatter plot (Figure 3h), the incorrect fit (blue) is partially obscured, while the correct fit (cyan) stands out in the upper-right corner. By integrating fitting probabilities and local z-scores, DomainSeeker correctly selected the optimal domain as the top candidate. This approach, which considers all possible fitting positions, effectively prevents missed identifications due to incorrect top-ranked fits.

Three densities (19, 21, and 26) were not correctly identified (Fig. S2c), reflecting two distinct factors (Fig. S10). Densities 19 and 21 were incompletely segmented: much of the correct domain lay outside the segmented region. For density 19, the captured fraction was too small to yield a meaningful fit.

Density 21’s correct placement was not a local correlation maximum and was absent from the candidate pool. Density 26 is intrinsically small: even minor segmentation imperfections leave too few features for reliable identification. Overall, domain–density mismatches are common in the results, yet only a few extreme cases led to identification failures, demonstrating that DomainSeeker is robust to moderate domain–density mismatches.

### Identifying domains using posterior probabilities

We examined the impact of XL-MS data on domain identification using the inner ring (IR, EMD-24232, 7.6 Å) and the outer ring (OR, EMD-24231, 11.6 Å) density maps of the yeast nuclear pore complex (NPC), whose corresponding atomic models are available (PDB entry 7N85 for IR, 7N84 for OR)^30^. In total, 31 structured proteins (Table S2) were selected, and density regions covering these proteins were manually segmented. Predicted structures of 5,300 candidate proteins from the yeast proteome in AFDB were parsed into 7,744 domains.

XL-MS data were incorporated into the calculation of posterior probabilities, with NPC symmetry explicitly considered. The dataset we used contained 2,167 distinct cross-links^41^; however, only 101 pairs corresponded to residue pairs within the cross-linking constraint (≤50 Å). This limited number mainly reflects the prevalence of self-cross-links and the fact that most cross-links involve flexible regions not included in the parsed domains. Of these, only 29 pairs contributed meaningfully to the analysis (Table S3).

In Figure 4a, cyan circles and green hexagons indicate rankings based on prior and posterior probabilities, respectively, while shaded areas indicate their probabilities. Using prior probabilities alone, DomainSeeker correctly identified 18 of 31 domains, with most missed cases corresponding to lower-resolution regions. Incorporating XL-MS data improved identification, revealing four additional domains (12, 24, 26, 27) and increasing fitting confidence for another four domains (1, 2, 11, 14). Detailed results are presented in Table S2. Structures and individual global and local results are shown in Figure S11.

**Figure 4.**
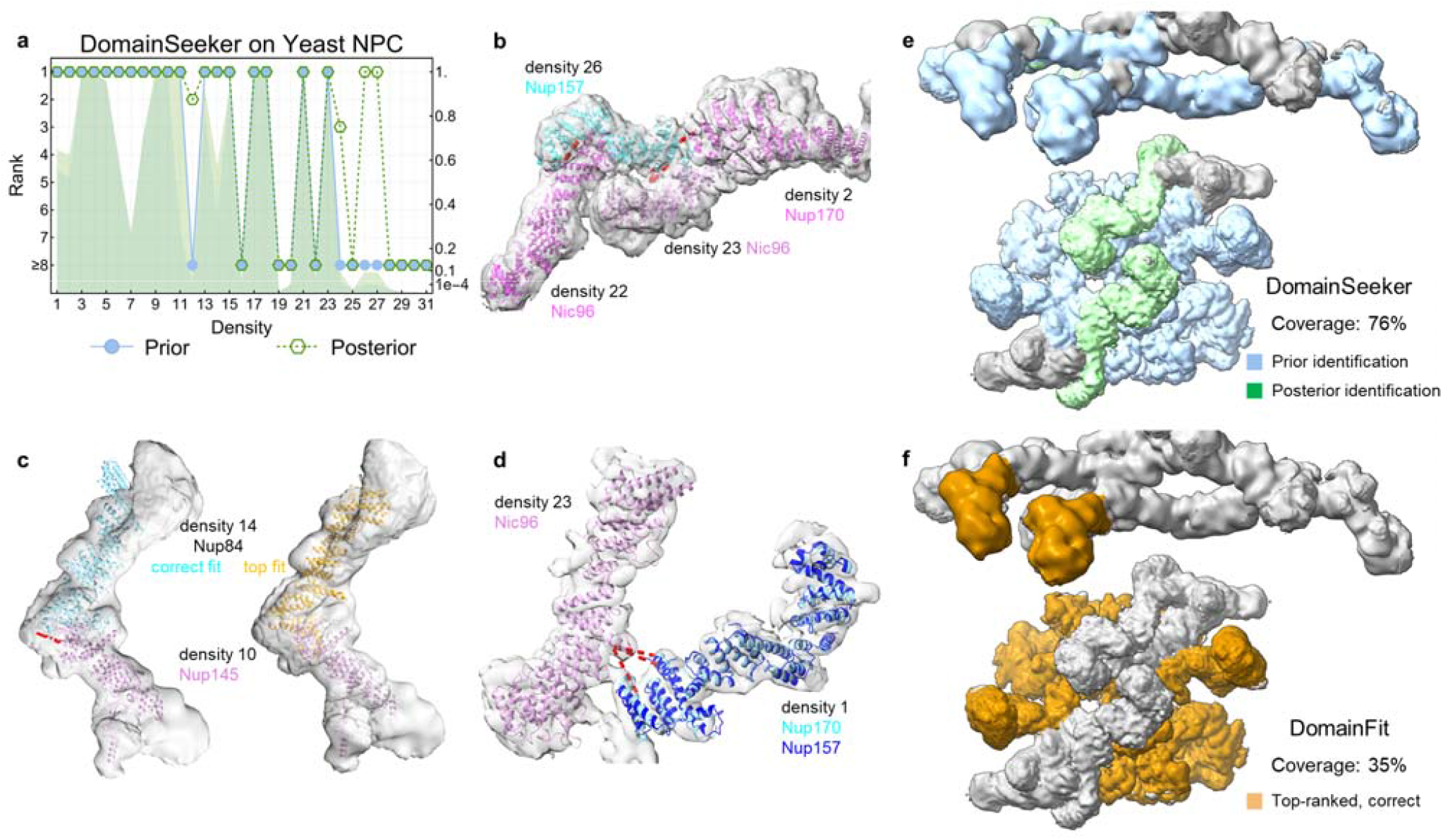
Performance on densities segmented from the yeast NPC. (a) Performance on the yeast NPC using prior (cyan) and posterior (green) probabilities. Circles/hexagons: rankings of the correctly fitted domains; shading: probabilities. (b) Nup157 fitted to density 26. Red dashed lines: cross-links between Nup157 (density 26) and the neighboring structures. (c) Nup145 correctly positioned in density 10 (pink); Nup84 in density 14 (cyan: the correct fit; orange: the top fit). Red dashed line: compliant cross-link. (d) Nup170/Nup157 fitted to density 1. Red dashed lines: cross-links between Nup170 (density 1) and Nic96 (density 23). (e) Regions identified by the results of DomainSeeker. Prior and posterior identification are displayed in cyan and green, respectively. The total coverage ratio is 76%. (f) Regions identified by the top-ranked results of DomainFit. The coverage ratio is 35%.

Notably, in Figure 4a, the correct proteins for densities 26 and 27 were unidentifiable using priors alone. After integrating XL-MS data, their rankings increased dramatically, placing them at the top of the posterior results. As shown in Figure 4b, Nup157 (density 26) likely forms cross-links with proteins in adjacent densities, substantially boosting both its posterior probability and ranking.

In Figure 4c, DomainSeeker correctly identifies the placement of Nup84 (cyan) within density 14, whereas ChimeraX ranks an incorrect fit (orange) as top. Structural prediction errors cause slight deviations in the upper terminal region of Nup84, leading ChimeraX to mis-rank the correct fit. In contrast, DomainSeeker systematically evaluates all candidates and successfully assigns the correct position as optimal. Although the prior probability of this fit ranks first at 35%—not exceptionally high—the presence of a cross-link between Nup84 and Nup145 (fitted in density 10, pink) raises its posterior probability to 56%, thereby increasing confidence in the correct assignment.

XL-MS data also assist in distinguishing homologous proteins. In Figure 4d, both Nup170 (cyan) and Nup157 (blue) fit well into density 1 and exhibit highly similar structures, resulting in comparable prior probabilities (55% and 45%, respectively). Incorporating cross-links between Nup170 (density 1) and Nic96 (density 23) increases the posterior probability of Nup170 to 65%, resolving the ambiguity.

Figures 4e–f compare the regions identified by DomainSeeker and DomainFit, which achieve coverage ratios of 76% and 35%, respectively. These results demonstrate that DomainSeeker delivers substantially higher identification accuracy.

Overall, posterior improvements consistently relied on a few cross-links bridging to reliably assigned partners. Subsampling confirmed this: at 10% of the original list, a single informative cross-link maintained the posterior near the full-set level (Table S4, density 14). What matters is not the total number, but a few key informative ones—those connecting two rigid regions where at least one corresponding density has sufficient resolution and completeness for reliable prior identification.

### Building preliminary structural models using a ChimeraX plugin

We further developed an interactive ChimeraX plugin for DomainSeeker (Figure 5). The plugin provides input fields for key parameters of the four workflow modules, with pre-set default values for convenience. Through this visual interface, users can readily run DomainSeeker to perform domain identification and directly view computational results, densities, and fitted structures within ChimeraX. Experimental data consistent with the models are simultaneously mapped—for example, cross-links are displayed as red dashed lines—facilitating intuitive inspection and filtering of results.

**Figure 5.**
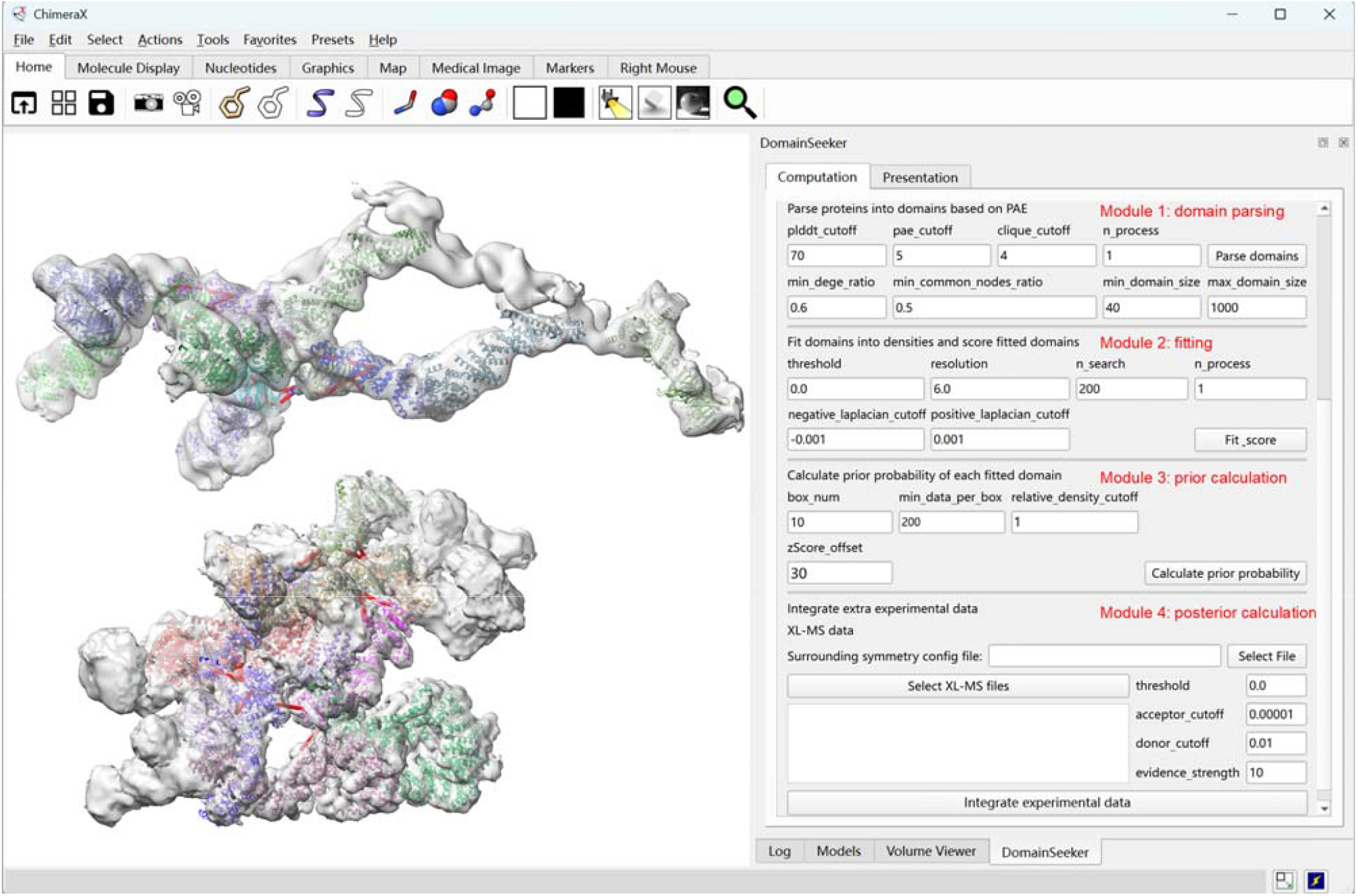
The preliminary model built by DomainSeeker GUI in ChimeraX. Left panel: the preliminary model built by DomainSeeker with cross-links denoted as red lines. Right panel: the graphical user interface of DomainSeeker in ChimeraX, facilitating the calculation of the four modules.

Using this plugin, we examined the results from the yeast NPC density maps to construct a preliminary structural model (Figure 5). For each density region, candidate states with probabilities above a threshold (1% in this study) were inspected. Regions lacking states above the threshold were considered unrecognized. By default, the top-ranked state was selected, although alternative states could be chosen when additional evidence, such as spatial conflicts, was observed. Following these criteria, all 22 domains included in the preliminary model were correct. This validates that DomainSeeker has strong potential to address the challenges in protein domain identification and modeling, especially for in situ cryo-ET.

## Discussion

In this study, we present DomainSeeker, an efficient computational workflow and software tool for identifying protein domains in cryo-ET density maps by integrating AlphaFold2-predicted structures with complementary experimental data. A dedicated ChimeraX plugin was developed to implement the workflow and streamline visualization and analysis.

DomainSeeker’s performance using prior probabilities alone was evaluated in the mouse sperm microtubule doublet. From the 6.5 Å cryo-ET map, 26 densities corresponding to known MIPs were segmented. DomainSeeker correctly identified all 26 domains, maintaining high accuracy even at 10 Å resolution, and successfully resolved an additional unmodeled density. When tested on 20 automatically segmented densities to emulate real-world in situ conditions, 17 domains were correctly identified. Across all tests, DomainSeeker significantly outperformed the existing algorithms in both accuracy and robustness across resolutions.

We next assessed DomainSeeker’s capacity to integrate XL-MS data using the yeast NPC maps at 7.6–11.6 Å resolution. Among 31 segmented densities, 18 domains were identified using prior probabilities alone, and incorporation of XL-MS data enabled the identification of four additional domains. The inclusion of XL-MS information improved recognition confidence, corrected misassigned fits, and distinguished homologous proteins. A preliminary structural model of the yeast NPC was then assembled, demonstrating that the ChimeraX plugin effectively supports model inspection and refinement.

Several key algorithmic advances underlie these improvements. It has been observed that residue networks derived from AlphaFold2 PAE matrices contain abundant clique structures—groups of residues with reliably predicted relative positions. Building on this, we developed a PAE-based domain parsing algorithm that isolates well-structured regions while filtering out flexible loops, improving fitting precision.

To evaluate domain–density matches comprehensively, a dual-level assessment combining global and local fitting metrics was designed. The local evaluation is based on a z-score derived from overlap volume and correlation, quantifying how strongly each fitted domain stands out relative to others with similar overlap volumes. As the number of fitted domains increases, these z-score distributions stabilize, providing statistically reliable estimates of local fitting quality.

Furthermore, we developed a workflow within a probabilistic framework to rationally incorporate XL-MS data. A key goal is to clarify how XL-MS data inform judgments about complex components and structures. Conversely, determining cross-links from a known biomolecular structure is straightforward. Bayesian principles clarify the relationship between these two problems. Through probabilization and Bayesian inference, we have derived the core mechanism by which XL-MS data support domain identification: summing contributions from the same cross-link pair across positions and multiplying those from different pairs.

We also confirmed that identification remains stable across reasonable ranges of the main parameters. Besides, a proteome-scale search confirmed that the results are not biased by candidate list composition (Fig. S5), and the test domains span broad ranges of molecular weight, shape, and local resolution (Fig. S12), together supporting the general applicability of DomainSeeker.

Moreover, DomainSeeker is not limited to monomeric structures; it can also directly utilize multimer-predicted complexes. When applied to AF3-predicted complexes^16^ of the yeast NPC, DomainSeeker identified cross-protein domains and successfully recognized them from their corresponding densities, in some cases outperforming monomer-level domain searches (Fig. S13). This highlights DomainSeeker’s flexibility to operate both as a standalone tool and as an enhancer within multimer-centric workflows.

Despite these advances, several challenges remain. First, the pipeline relies on the accuracy of predicted structures; prior probabilities alone cannot reliably distinguish homologs with highly similar conformations, requiring supplementary experimental data. Second, our pipeline has not yet automated density segmentation. At medium to low resolutions, accurately segmenting densities of non-spherical proteins is challenging, and individual domains often exhibit lower signal-to-noise ratios. Both factors increase the difficulty of reliable recognition. In addition, while considering all plausible fitting positions reduces omissions due to incorrect top-ranked fits, missing the correct position in the candidate list remains a limitation that requires improved fitting algorithms.

A core strength of DomainSeeker is its flexible probabilistic framework, which can integrate diverse experimental datasets. While we have initially incorporated MS and XL-MS data, future expansions could include additional modalities. For example, immunoelectron microscopy^42^, super-resolution microscopy^43^, and spatial mass spectrometry^44^ provide protein spatial localization that could refine prior probabilities when combined with MS-derived candidate lists. Meanwhile, co-immunoprecipitation^45^, proximity labeling^46^, and protein-protein interaction networks incorporating experimental or predictive data^45,47^ may offer interaction information to calculate posterior probabilities. Overall, given its considerable potential for further development, DomainSeeker is well-positioned to significantly advance research in in situ structural biology.

## Methods

### Segmentation

Densities of interest can be segmented from the full map either manually or automatically in ChimeraX. Users can place markers on regions that may correspond to potential domains, use the Color Zone tool to color the regions around these markers and the *volume splitbyzone* command to segment the densities. Alternatively, the Segment Map tool^48^ can be used for automatic map segmentation.

### Domain parsing

The AlphaFold2 predicted models and PAE files can be downloaded from AFDB using the *fetch_pdb_pae*.*py* script with an input list of UniProt IDs for the candidate proteins. Then, DomainSeeker provides a Python script *parse_with_pae*.*py* to divide the candidate proteins into domains. For domain parsing, the current default parameters achieve satisfactory performance: pLDDT_cutoff = 70, PAE_cutoff = 5, clique_cutoff = 4, min_edge_ratio = 0.6, min_common_nodes_ratio = 0.5, min_domain_size = 40, and max_domain_size = 1000.

### Fitting

The *fit_with_chimerax*.*py* script in DomainSeeker uses the *fitmap* command in ChimeraX to identify all potential fitting positions, recording correlation and hit metrics for global assessment, as well as overlap volume and correlation metrics for local assessment.

ChimeraX finds candidate fittings by performing gradient ascent optimization from a number of randomly generated initial search positions. For the test set of this work, an initial search number of 200 in ChimeraX is sufficient to find the correct fitting position and orientation. Users need to adjust this number along with the other parameters (threshold, resolution, negative_laplacian_cutoff, and positive_laplacian_cutoff) according to the input density map.

### Prior probability calculation

The *calculate_prior_probabilites*.*py* script calculates prior probabilities according to the cross-correlation and hits of every fitted domain, together with the overlap volume and correlation.

For a given density *A* and domain *a*, the probability that fitting α is the correct fitting scales proportionally with the product of the correlation-derived probability density and its associated hits

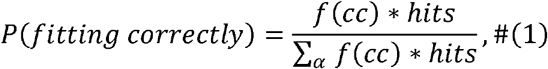

where *cc* represents the cross-correlation and *hits* represents the population of maxima within the cluster given by ChimeraX. The denominator represents the summation over all fitting positions of domain *a* into density *A* reported by the *fit_with_chimerax*.*py* script.

The function converting cross-correlation into probability density defined in DomainSeeker is

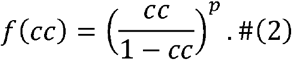

The transformation maps cross-correlation in [0, 1) to a positive unbounded evidence scale [0, +∞), satisfying the constraints that a perfect correlation yields overwhelming evidence while zero correlation yields none. The exponent p controls the nonlinearity of this mapping. Within our validation scope, all exponents yield similar and satisfactory results (Fig. S14a). We selected p = 3, which maintains robust ranking performance while avoiding potential excessive noise amplification.

The expression of the local z-score is

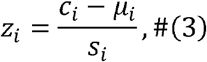

where *c*_*i*_ is the overlap correlation of the candidate fitted domain *i, μ*_*i*_ is the average score of the fitted domains with similar overlapping volume, and *s*_*i*_ is the corresponding standard deviation.

The prior probability is calculated according to

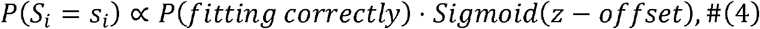

where *z* is the local z-score. The *offset* is used to prevent extreme dominance of the prior probability by excessively large z-scores, which may arise from insufficient data or other numerical artifacts.

In the script, the parameters—box_num, min_data_per_box, and relative_density_cutoff—are used to discretize the overlapping volume into intervals to estimate local z-scores.

### Integration of XL-MS datasets

DomainSeeker further enables the integration of prior probabilities with supplementary experimental datasets to infer posterior probabilities through Bayesian inference.

From a methodological perspective, it is imperative that supplementary experimental datasets influence the entire density map holistically rather than acting on isolated density regions, as such datasets inherently characterize the full biomolecular complex. Consequently, the probability that the entire system occupies a given global state should be considered.

The prior probability assumes independence among distributions for individual densities. Thus, the system probability is given by

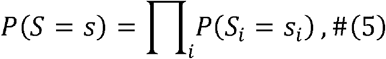

where *S* denotes a random vector representing the holistic structure across all target densities and *s* is a specific conformational instance of the full system. *S*_*i*_ and *s*_*i*_ correspond to density *i*. When considering extra experimental datasets, the posterior probability can be inferred by Bayesian inference, that is,

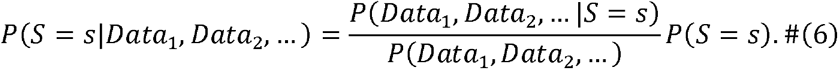

Datasets from different experiments should be independent, and thus, the posterior probability equals

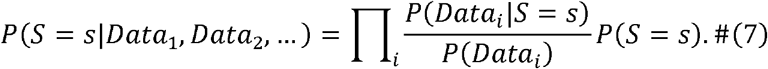

This equation implies that datasets from distinct experimental methods can be multiplied onto the prior probabilities as independent scaling factors.

To date, DomainSeeker has formalized scaling factors for XL-MS datasets. For a specific XL-MS dataset *CL*, the scaling factor represents the conditional probability of observing *CL* given a state *s*. It is calculated as the product of two probabilities: the probability of observing all cross-links in *CL* and the probability of detecting none of the structurally possible cross-links that are absent from *CL*.

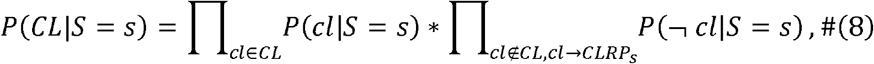

where *cl* represents a cross-link pair. Within the probability function’s argument parentheses, *cl* the event that the specific cross-link pair *cl* is observed, while ¬*cl* represents the complementary event. Given structure *s*, the set of crosslinkable residue pairs is denoted as *CLRP*_*s*_, while *cl*→*CLRP*_*s*_ means that *cl* is a potentially observable cross-link pair.

First, focus on the probability of detecting none of the structurally possible cross-links that are absent from *CL*. It is straightforward to demonstrate that this probability expands to:

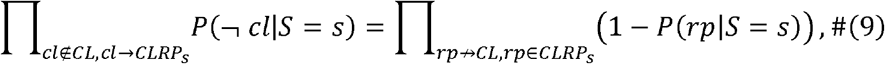

where *rp* represents a residue pair. *rp*↛*CL* means that the residue pair has no associated cross-link in the dataset. Within the argument set of the probability function, *rp* denotes the event that the residue pair is experimentally observed to form a cross-link. This equation represents the probability that all residue pairs cross-linkable in structure *s* yet absent from dataset *CL* are not detected.

Typically, the C_α_ -C_α_ distance distribution of cross-linked residue pairs follows a log-normal distribution^49^. Therefore, we modeled the probability of detecting a cross-link between a given residue pair as proportional to this log-normal distribution function of the C_α_ distance:

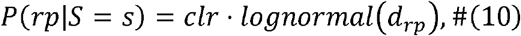

where *clr* is the cross-linking ratio, which represents the upper bound of the cross-linking probability for a residue pair. *d*_*rp*_ represents the C_α_ -C_α_ distance of the residue pair *rp*. The *lognormal* function is defined as a scaled version of the log-normal probability density function, with its maximum normalized to 1.

Considering that the ratio of the number of crosslinked residue pairs to the number of crosslinkable residue pairs is usually low^50^, we treat *clr* as a negligible quantity (*clr*≤1).

Under the small *clr* assumption, the second term of the scaling factor becomes negligible. While the probability of observing all cross-links in *CL* can be expanded to

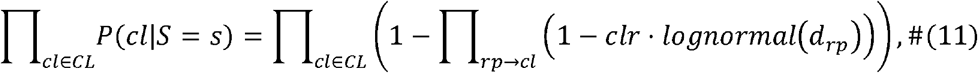

where *rp*→*cl* means the residue pair *rp* in the protein complex associates with the cross-link pair *cl*. Under the small *clr* assumption, this probability can also be simplified to

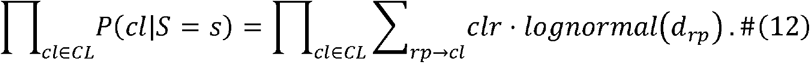

In fact, we have no need to assign a specific value to *clr*. It is apparent that *clr* appears homogeneously in both the numerator and denominator of the posterior probability expression, thus *clr* can be algebraically canceled out.

Given structure state *s*, residue pairs associated with a given cross-link *cl* are not necessarily localized in structure *s*. Therefore, the expression above can be rewritten as

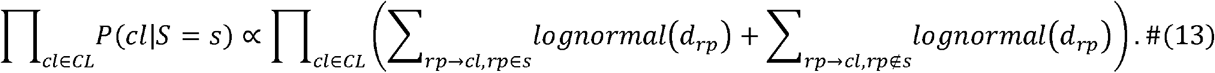

Actually, the latter term associated with the residue pairs outside structure *s* (outside the segmented density regions or excluded from the parsed domains) cannot be calculated due to insufficient information. Hence, we assign it an arbitrary numerical value ε as a heuristic approximation.

The above expression can then be rewritten as

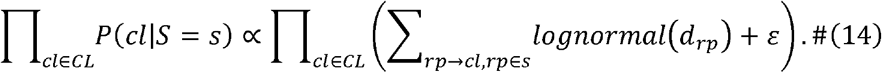

Moreover, for enhanced concision,

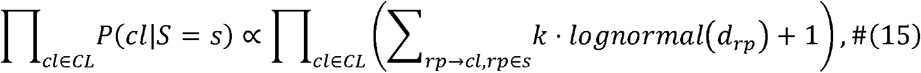

where *k*=1/ε. The parameter k weights the relative contribution of the XL-MS evidence in the posterior distribution. It is typically determined empirically or through iterative testing.

As derived above, the final posterior probability can be approximated using the following expression:

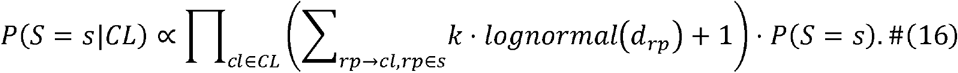

This expression indicates that the posterior probability of a given state is proportional to the prior probability multiplied by a scaling factor determined by the XL-MS dataset.

### Posterior probability calculation

The *calculate_posterior_probabilites*.*py* script in DomainSeeker can be executed to compute posterior probabilities incorporating XL-MS datasets.

We provide a simple symmetry-handling mechanism. Users specify the symmetry operations that map the initial unit cell to adjacent positions in a JSON file. Since biological macromolecular assemblies typically involve only rotational and translational symmetry, users only need to provide, for each relevant symmetry operation, the symmetry axis, a reference point, the rotation angle and the translation vector, together with the list of affected density regions. DomainSeeker then automatically generates symmetry-related copies and maps cross-link data onto them. For interactions mediated by symmetry, only the asymmetric interfaces need to be considered. For example, in the case of the C8 symmetry of the NPC, specifying a 45° rotation about the symmetry axis is sufficient; the −45° transformation is unnecessary because the corresponding interface is symmetrically equivalent and provides no additional information.

### DomainSeeker as a ChimeraX extension

The interface includes two tabs: one for computations and the other for interactive result visualization. In the computation tab, the plugin can automatically download predicted structures and PAE files, and users can also perform domain parsing via the plugin. Next, domain fitting is performed, followed by prior and posterior probability calculations. Once computations in the computation tab are completed, the results can be presented in the presentation tab, showing the domains, fit IDs, prior probabilities, prior rankings, posterior probabilities and posterior rankings for the selected state. In addition, ChimeraX visualizes all density maps and their fitted domains for the current state, with supporting evidence (such as cross-links) mapped on the structures.

### Computational resources

All computational experiments in this study were performed using ChimeraX 1.10 (embedded Python 3.11), with dependencies managed within the ChimeraX environment. DomainSeeker has been tested on Windows, Linux, and macOS with ChimeraX ≥ 1.6. The plugin can process approximately 4,000 candidate proteins across ~30 density regions within 1–2 days on a standard multi-core workstation (detailed benchmarks in Table S5).

## Data availability

The cryo-ET and cryo-EM density maps used in this study were obtained from the Electron Microscopy Data Bank under accession codes EMD-35230, EMD-35823, EMD-24232, and EMD-24231. The corresponding atomic models were retrieved from the Protein Data Bank under accession codes PDB 8I7R, 7N85, and 7N84. Mass spectrometry data, candidate protein lists, and XL-MS datasets are available in the supplementary materials of the respective studies. For the yeast NPC, candidate proteins comprised the complete *Saccharomyces cerevisiae* proteome retrieved from AFDB. AlphaFold2-predicted structures and PAE files for all candidate proteins, as well as TED domain annotations, were downloaded from AFDB. All other data are available from the corresponding authors upon reasonable request. Source Data are provided with this paper.

## Code availability

The code for DomainSeeker is available at https://github.com/zyzhangGroup/DomainSeeker.

## Acknowledgments

This work is supported by the National Key Research and Development Program of China (2021YFA1301504 to Z.Z., 2024YFA1307402 to Y.Z.), the Chinese Academy of Sciences Strategic Priority Research Program (XDB37040202 to Z.Z.), the National Natural Science Foundation of China (32525005 to F.S.), the Beijing Natural Science Foundation (JQ24056 to Y.Z.), the Basic Research Program Based on Major Scientific Infrastructures, CAS (JZHKYPT-2021-05 to F.S.), and the Fundamental and Interdisciplinary Disciplines Breakthrough Plan of the Ministry of Education of China (JYB2025XDXM502 to Z.Z.). The Supercomputing Center of USTC provides partial computer resources for this project. We are grateful to Mr. Yundong Zhang for technical support.

