## Supplementary Figures 1-14 for "De novo identification of protein domains in cryo-electron tomography maps from AlphaFold2 models"

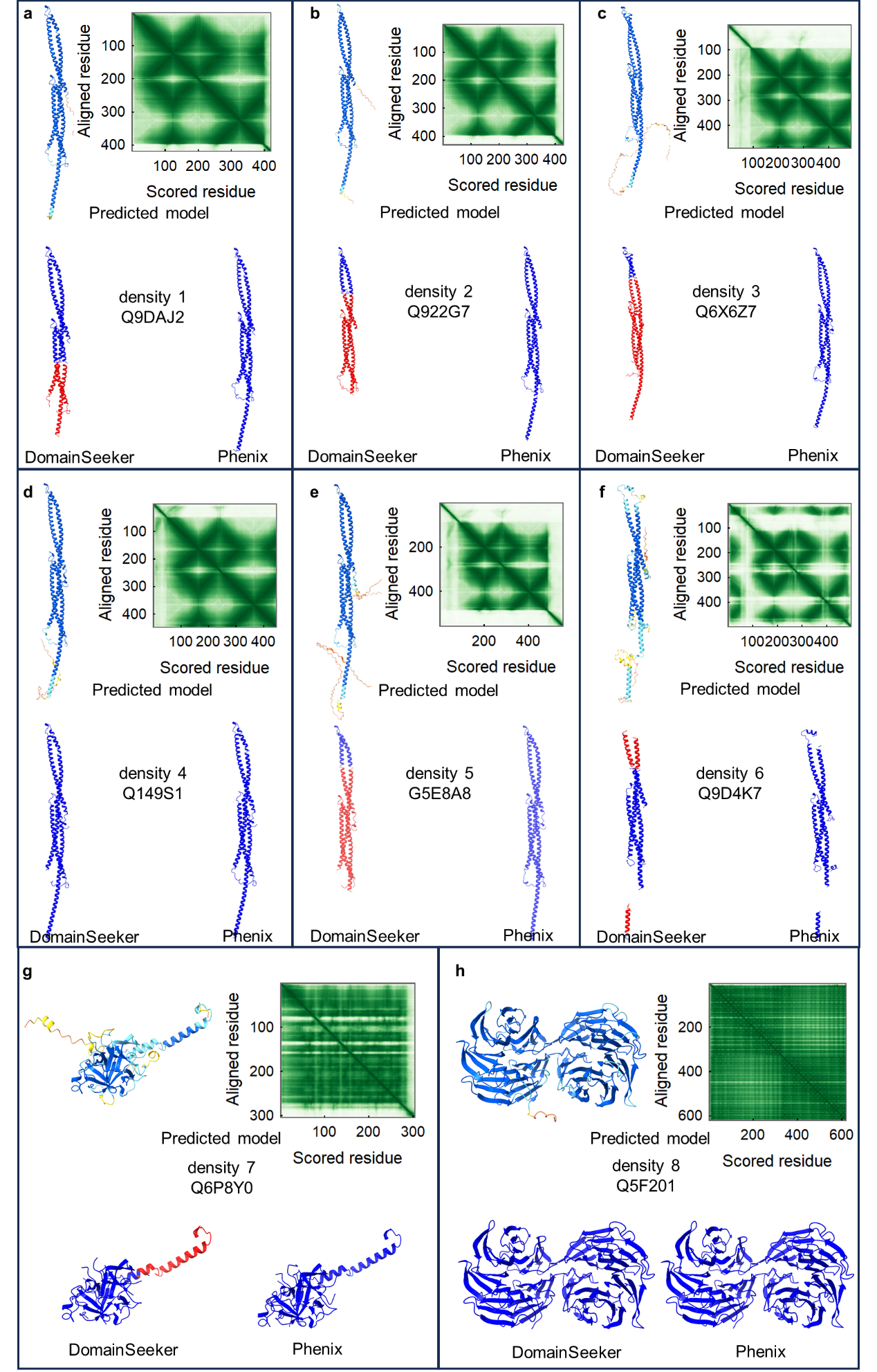

**(Continued on the next page)**

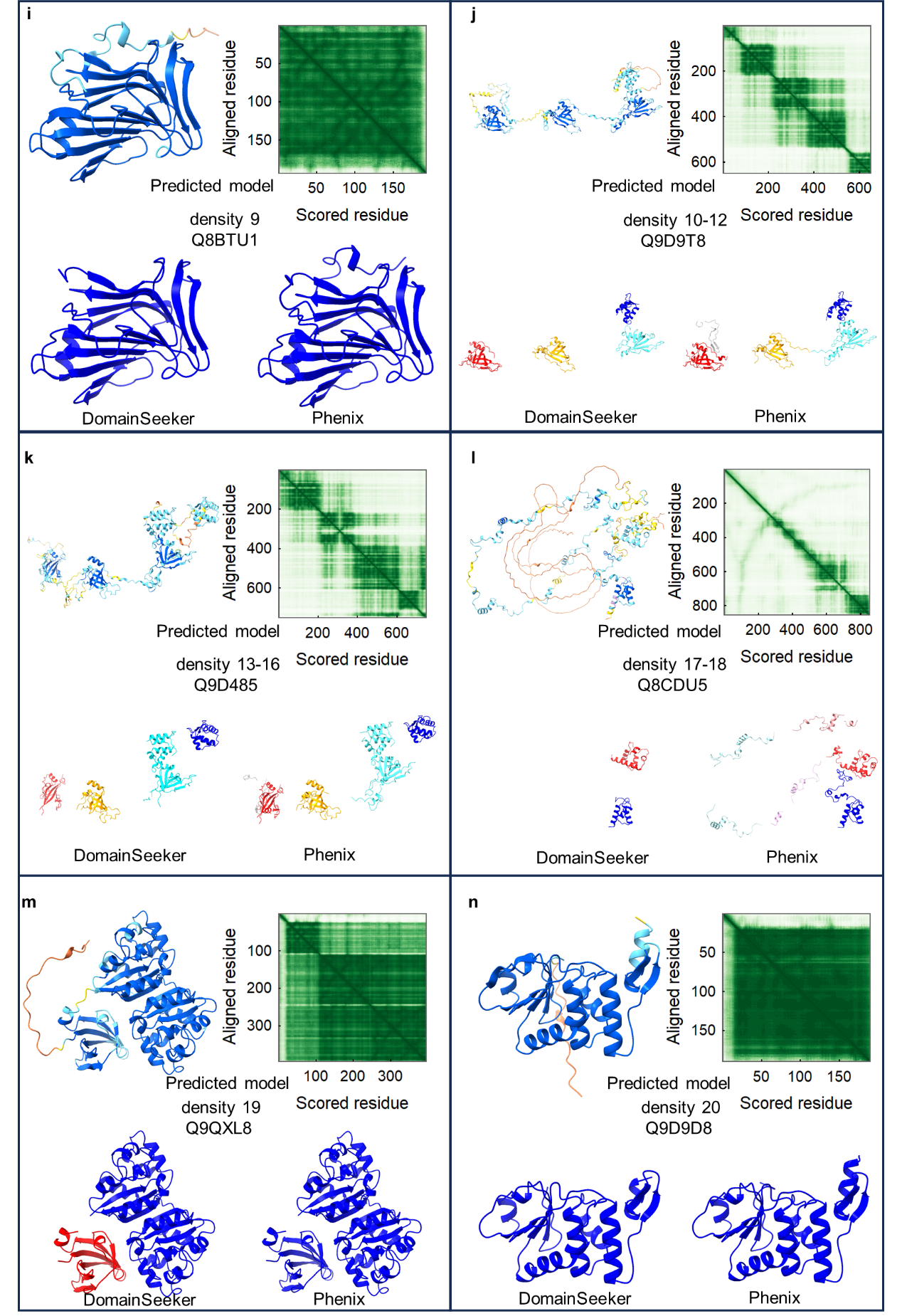

**(Continued on the next page)**

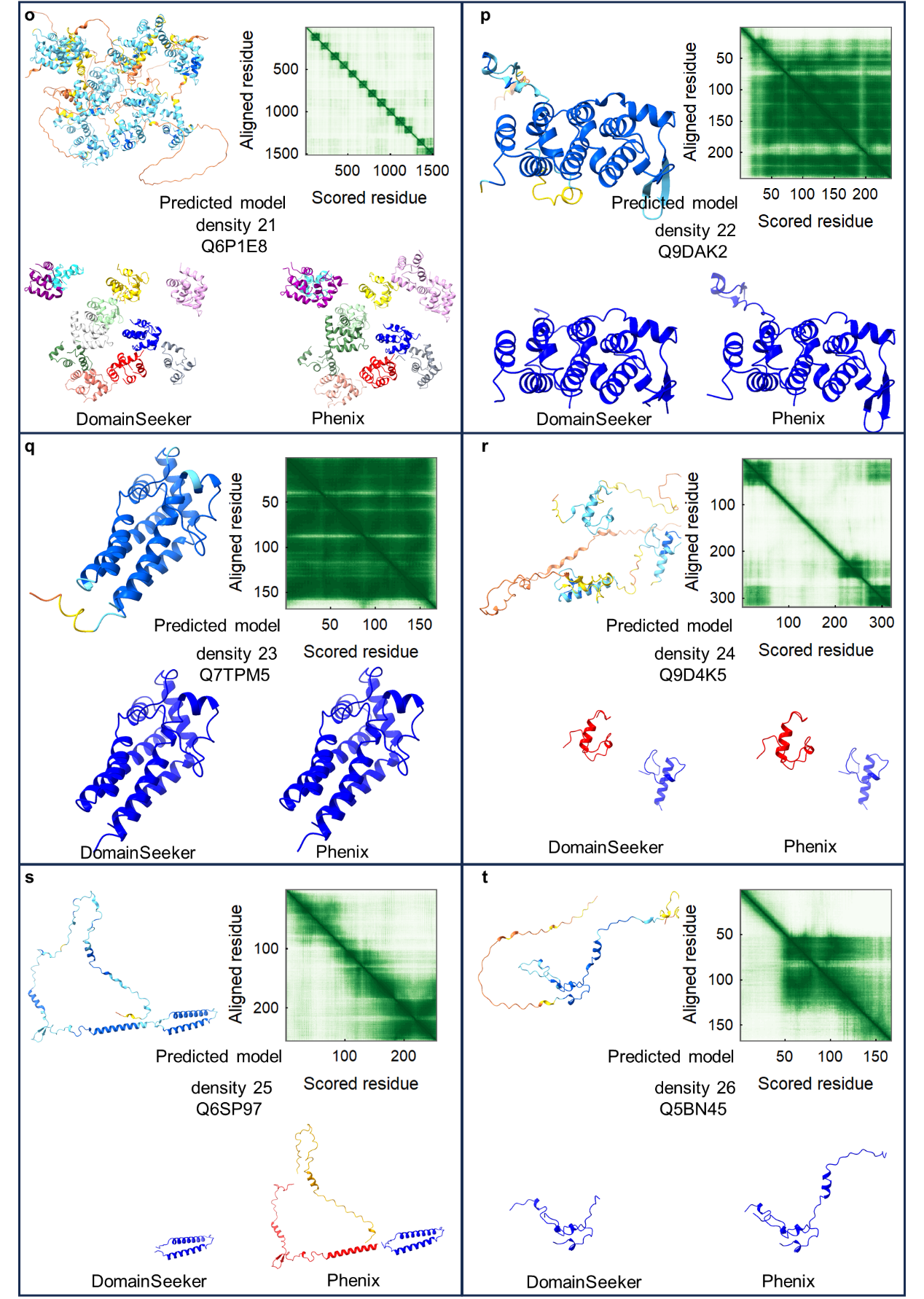

**Figure S1. Domains** **parsed by DomainSeeker and Phenix**

Domains parsed by DomainSeeker and Phenix (with PAE) in 20 proteins of the mouse sperm microtubule doublet. In each panel, the upper-left displays the AF2 predicted model colored by its pLDDT score. The upper-right shows the PAE graph, where darker shades of green indicate smaller PAE values. The lower-left and lower-right illustrate domains parsed by DomainSeeker and Phenix, respectively, with different domains represented by different colors.

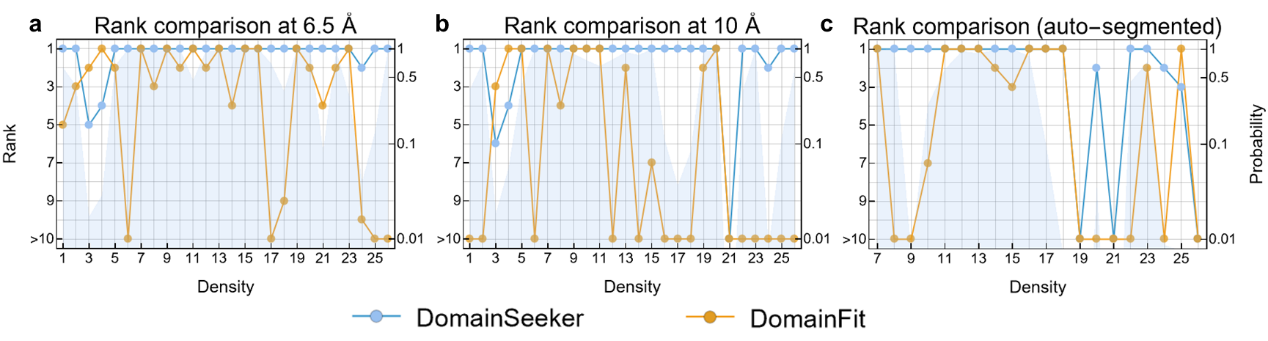
**Figure S2. Results on densities segmented from the mouse sperm microtubule doublet**

(a-c) Performance of DomainSeeker (cyan) and DomainFit (orange) on model-based densities (6.5 Å and 10 Å) and auto-segmented densities (6.5 Å). Circles: rankings of the correct domain fittings; cyan shading: prior probabilities given by DomainSeeker.

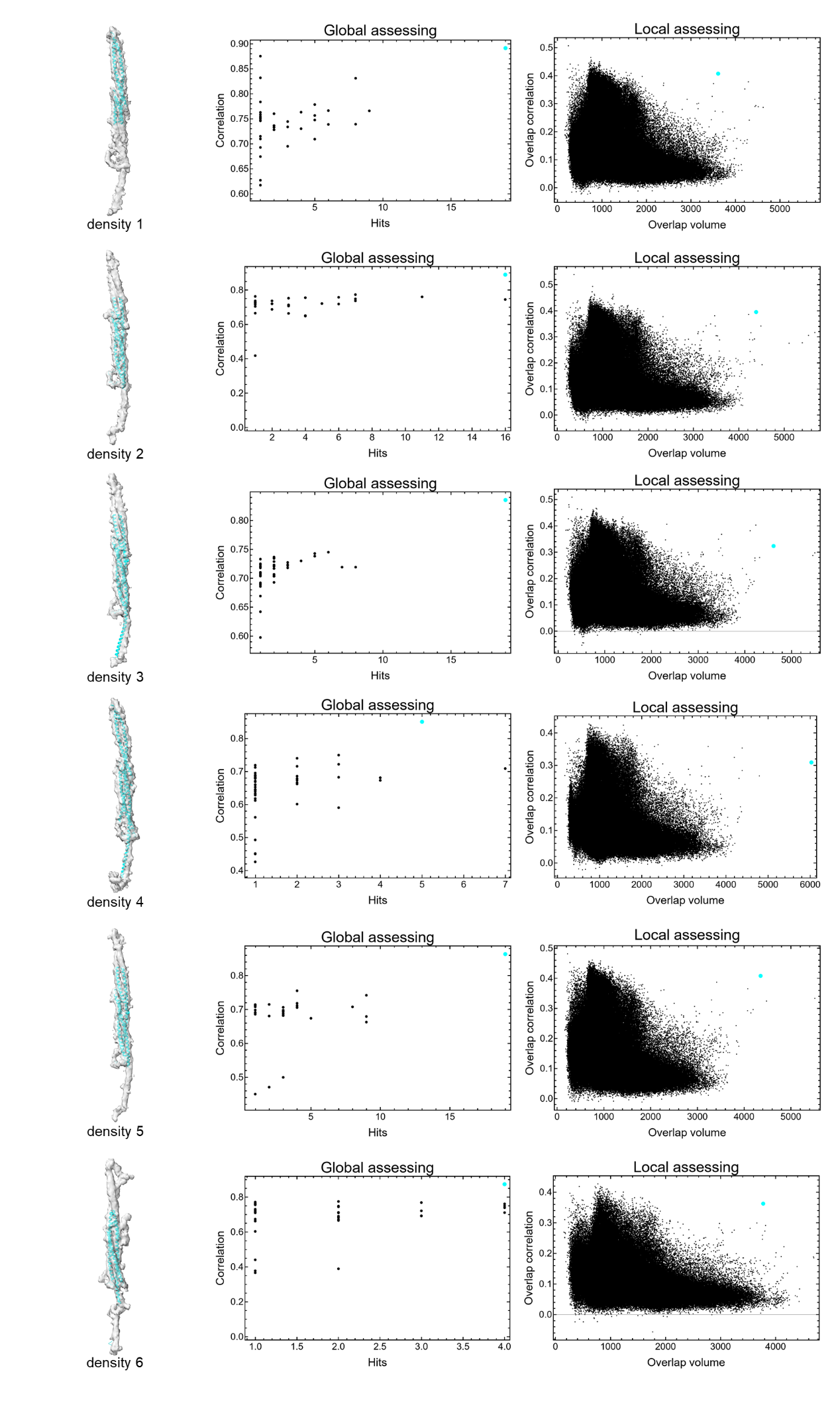

**(Continued on the next page)**

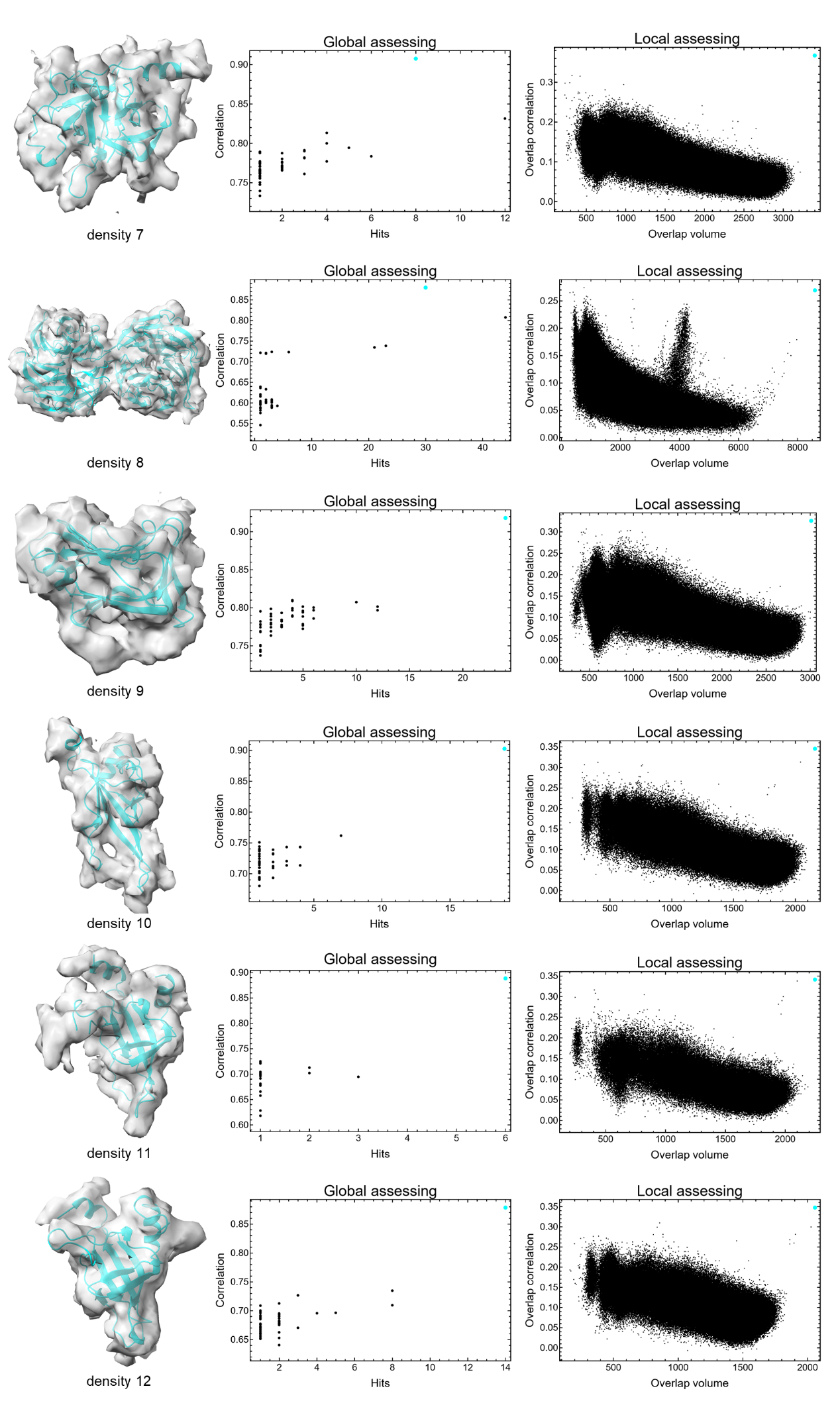
**(Continued on the next page)**

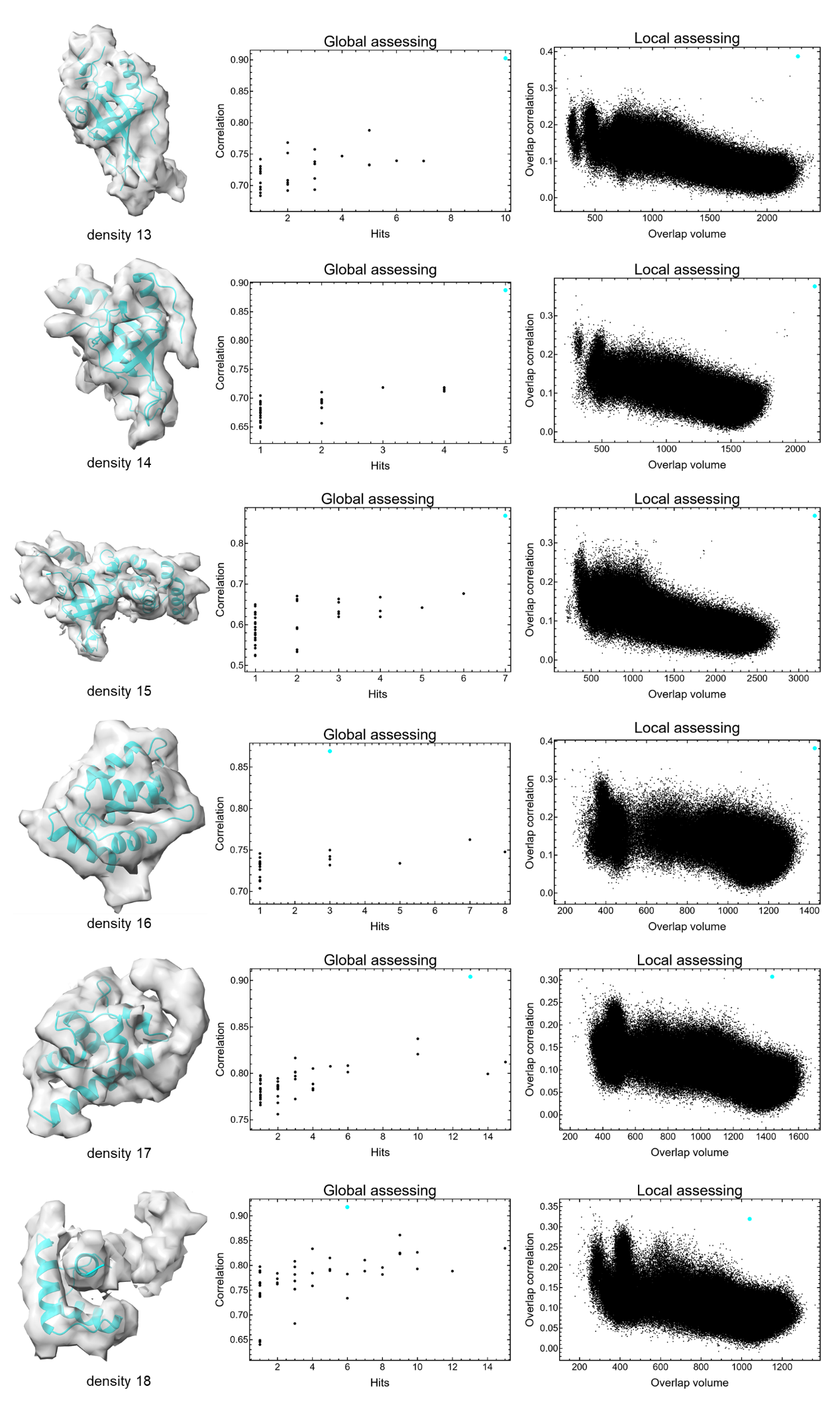
**(Continued on the next page)**

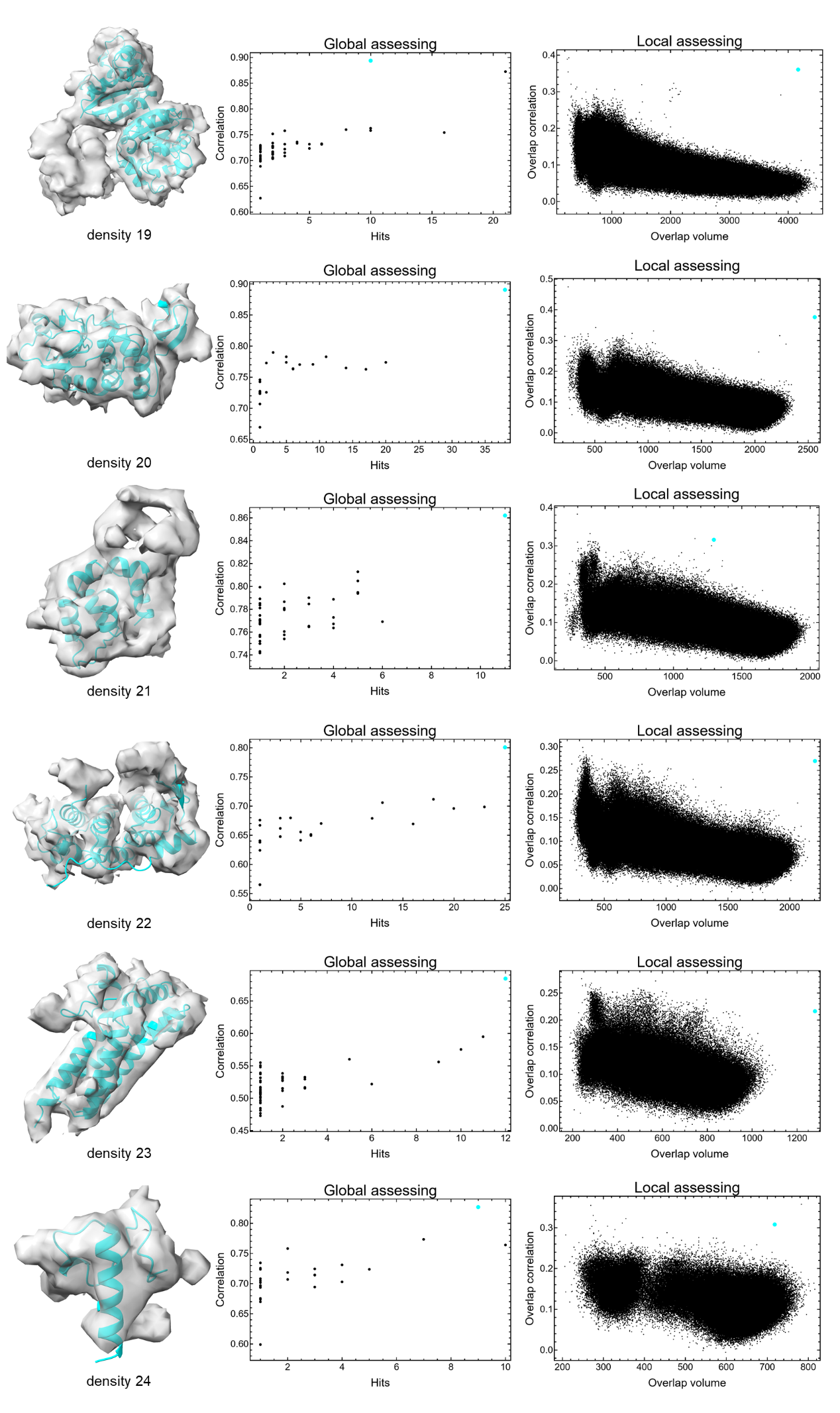
**(Continued on the next page)**

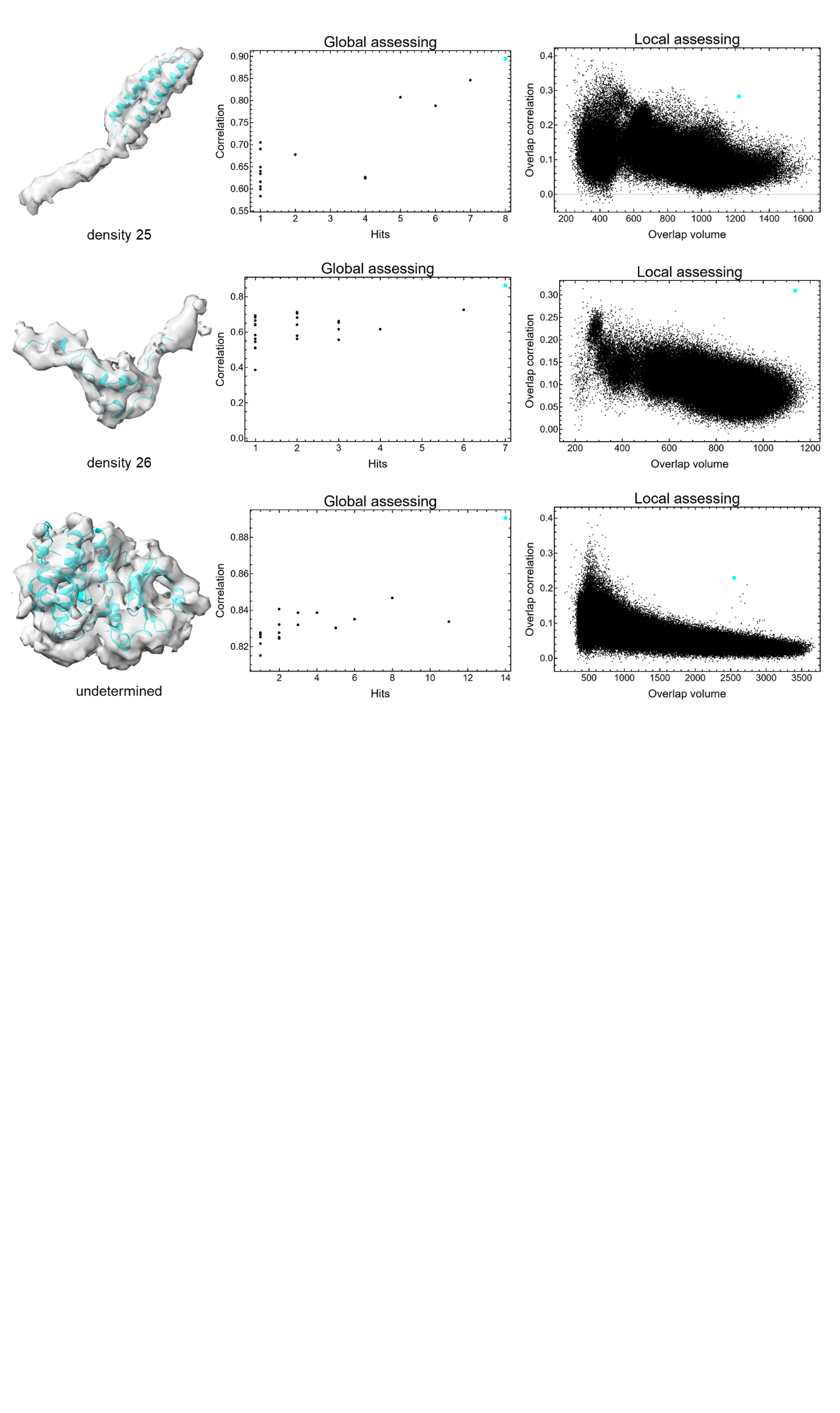

**Figure S3. Intermediate results of DomainSeeker on model-based densities at 6.5** **Å from the mouse sperm microtubule doublet**

Each row represents a density. The left panel: the correct fitting position of the correct domain (cyan structure). The middle panel: hits and correlation values of candidate fittings for the correct domain (the cyan point indicates the correct position). The right panel: overlap volumes and correlations of all domain fittings (the cyan point marks the correct fit).

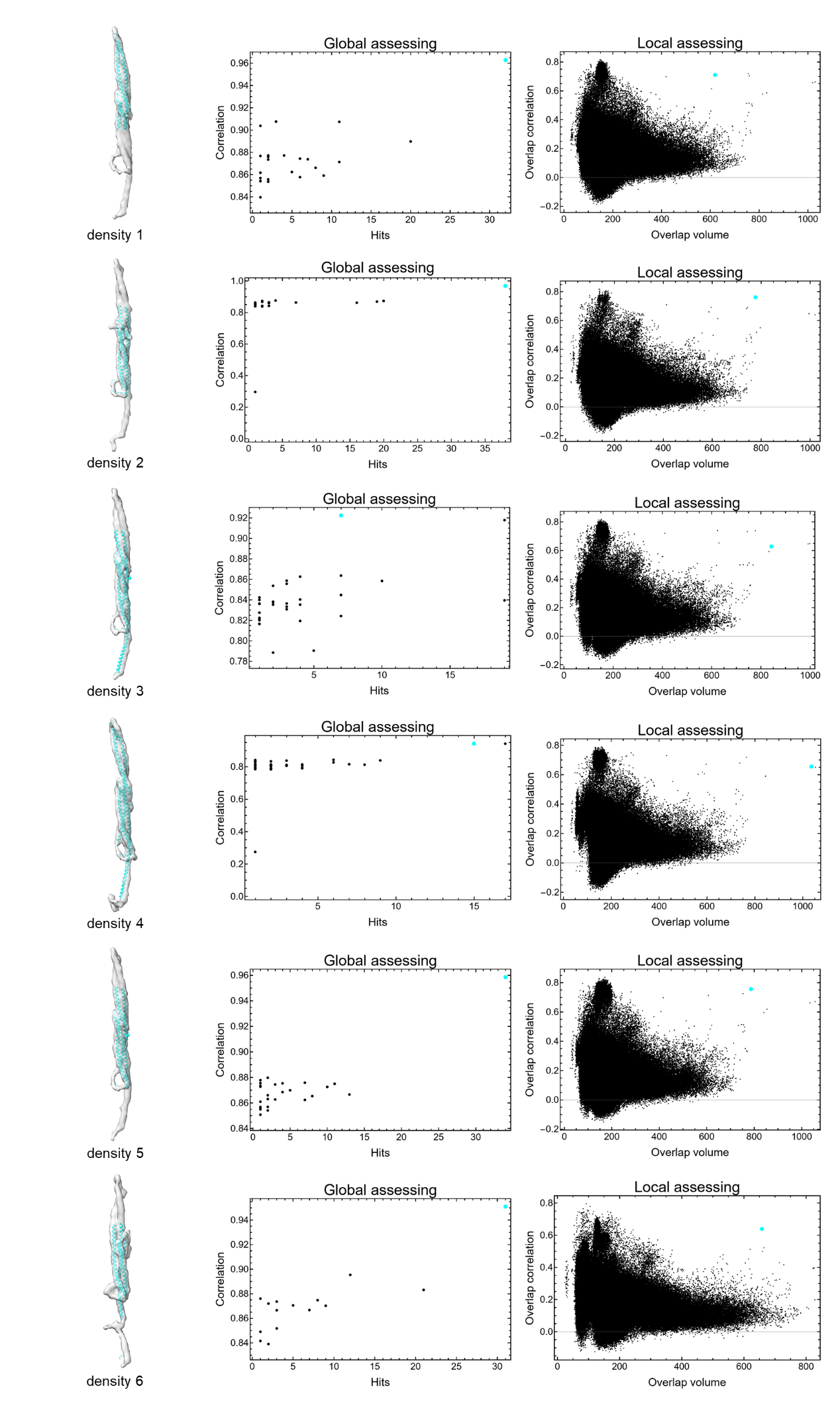
**(Continued on the next page)**

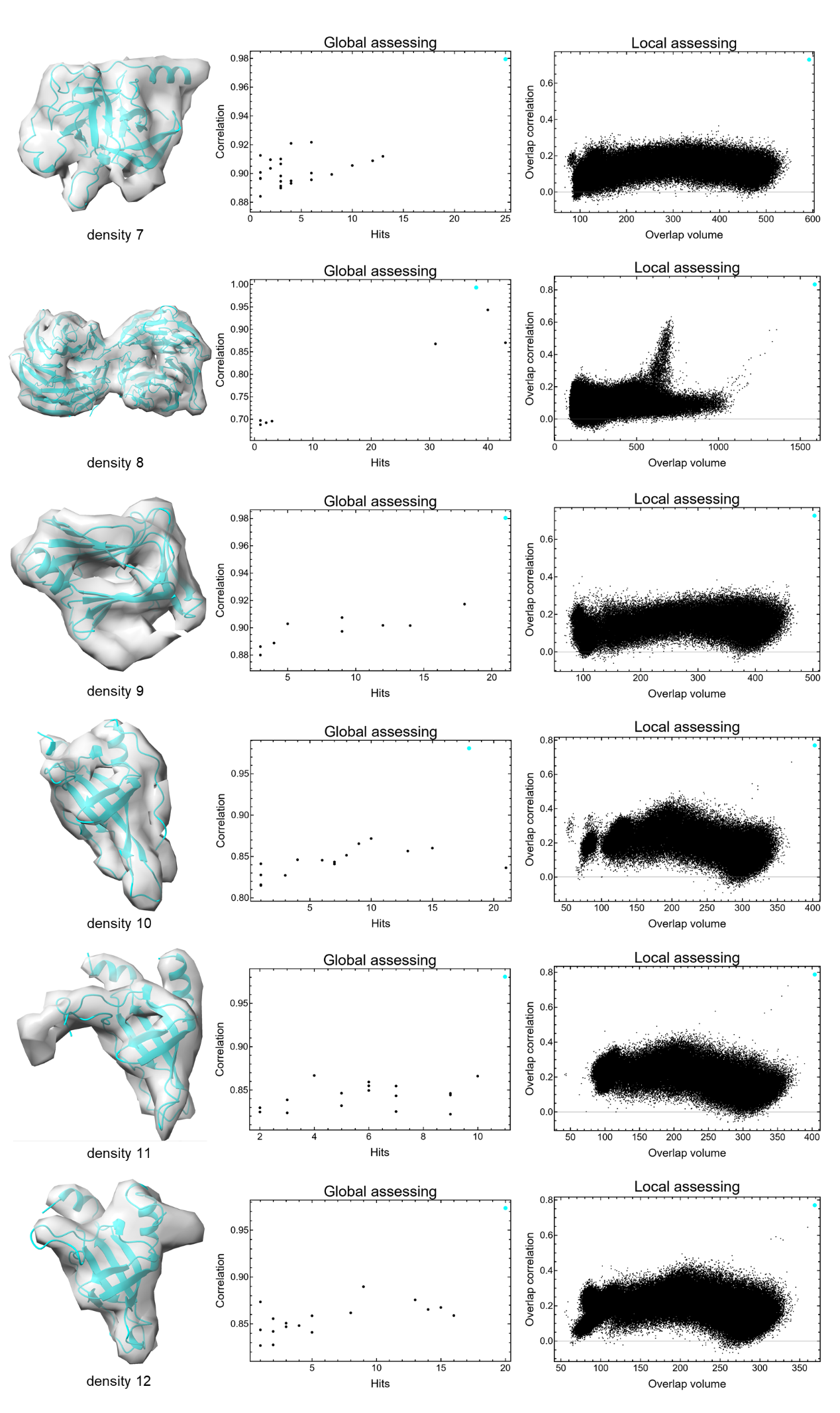
**(Continued on the next page)**

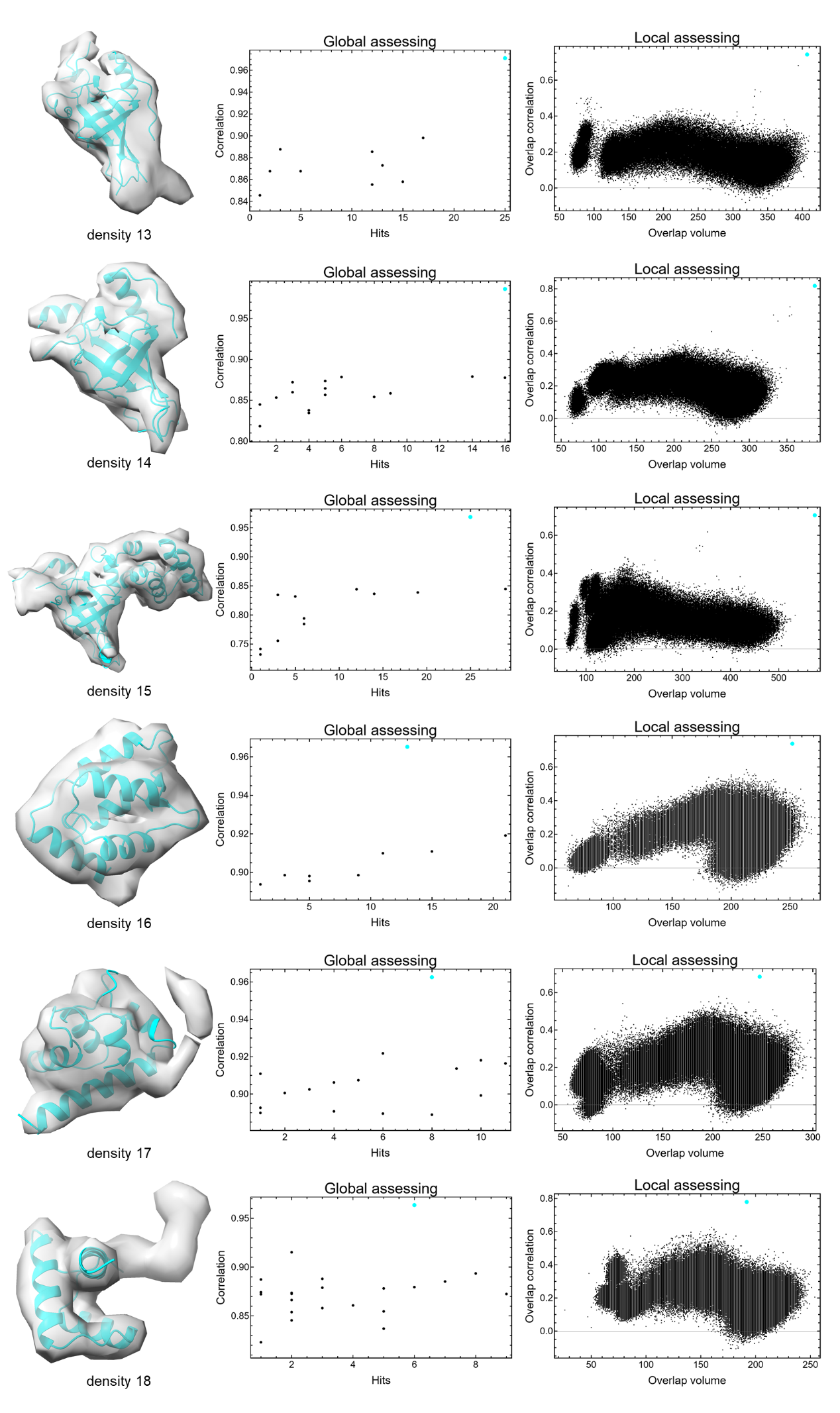
**(Continued on the next page)**

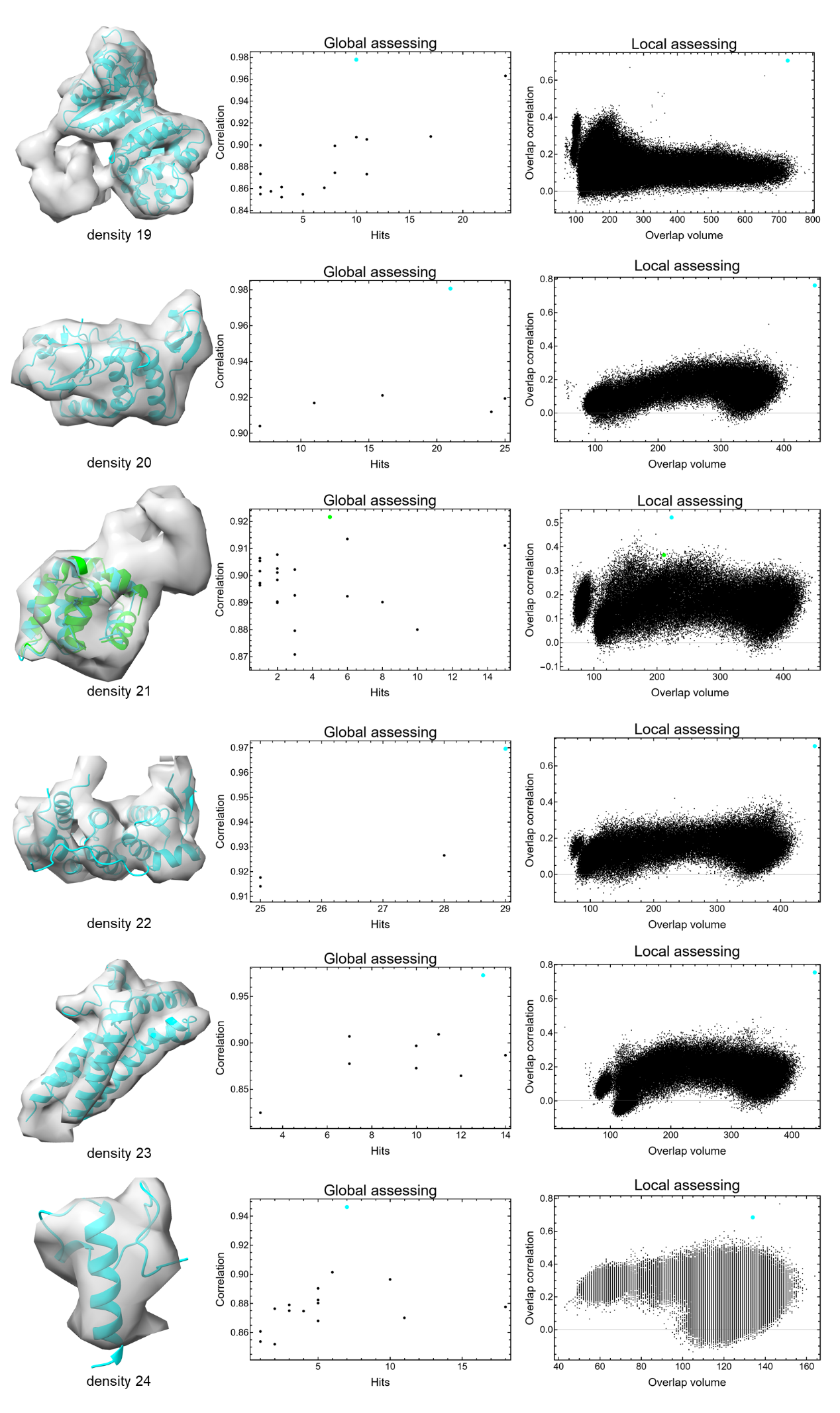
**(Continued on the next page)**

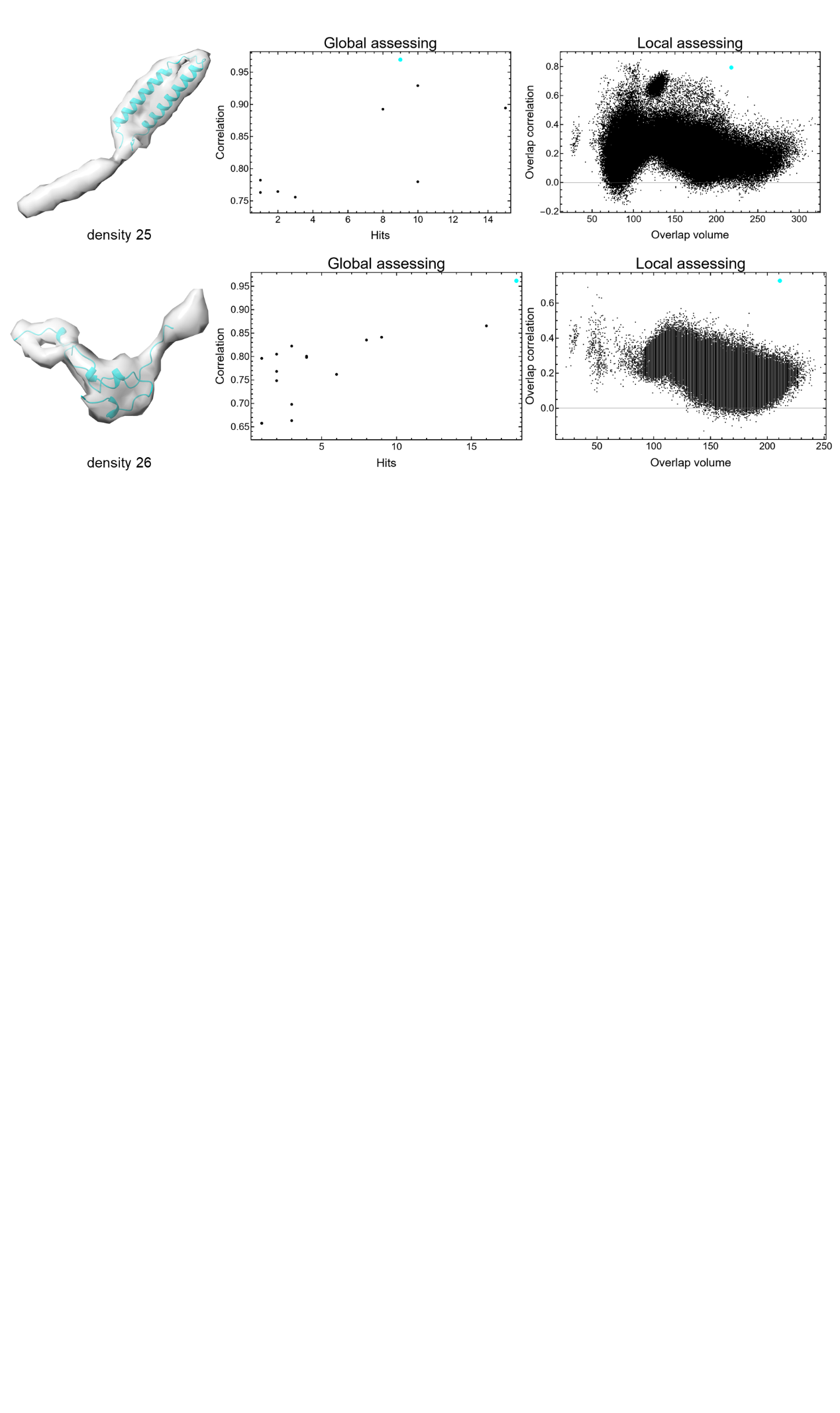

**Figure S4. Intermediate results of DomainSeeker on model-based densities at 10** **Å from the mouse sperm microtubule doublet**

Each row represents a density. The left panel: the correct fitting position of the correct domain (cyan structure). The middle panel: hits and correlation values of candidate fittings for the correct domain (the cyan point indicates the correct position). The right panel: overlap volumes and correlations of all domain fittings (the cyan point marks the correct fit). For density 21, the cyan point (the correct fit, yet not the local correlation maxima) in local assessment plot is manually added (not in the raw data); The green point/structure denotes the local maxima.

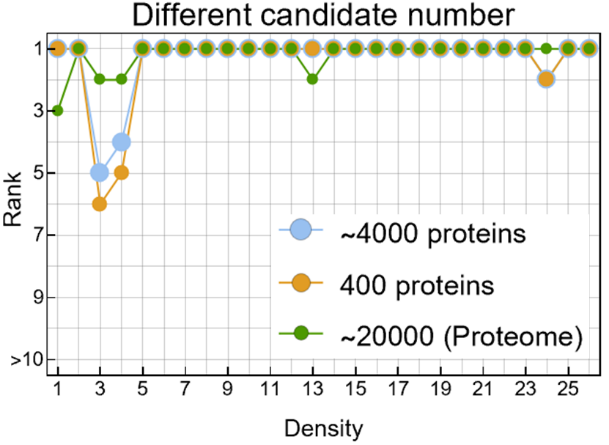

**Figure S5. Identification accuracy is robust to candidate set size and composition**

Domain identification rankings on the model-based 6.5 Å sperm MIP densities were obtained using three candidate sets: the full MS-derived set of over 4,000 proteins (cyan), a reduced set of 400 proteins including all ground-truth proteins (orange), and the complete mouse proteome from AFDB of approximately 20,000 proteins (green). The rankings are nearly identical across all the three sets, with all correct domains identified at high ranks. Minor differences in z-score estimates become more stable as the candidate set size increases. The large and compositionally diverse proteome-scale set further confirms that DomainSeeker's performance is not biased toward a specific candidate set composition.

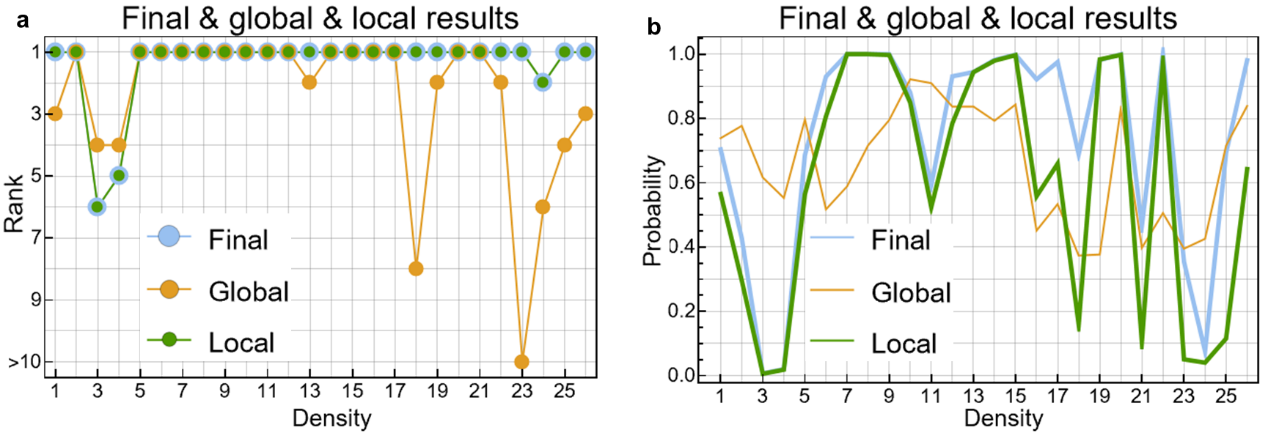

**Figure S6. Ablation analysis of global and local scoring components**

Recognition accuracy on the model-based 6.5 Å sperm MIP dataset under three scoring settings: global cross-correlation only (orange), local z-score only (green), and the combined scoring function (cyan) used in DomainSeeker. (a) Ranks: ranks of the local results are the same as those of the final results. Global results also achieve fairly high overall identification accuracy, even outperform the final results on densities 3 and 4, but are less stable. (b) Probabilities: The global probability is normalized across different fitting positions of the same domain. Final probabilities are elevated relative to local results across the board, owing to the complementary global similarity provided by the global correlation. This elevation in probability may, in certain cases, translate into a meaningful improvement in ranking. Together, the two metrics achieve complementary robustness: the local z-score ensures strong discriminative power across most cases, and the global correlation safeguards against localized structural mismatches.

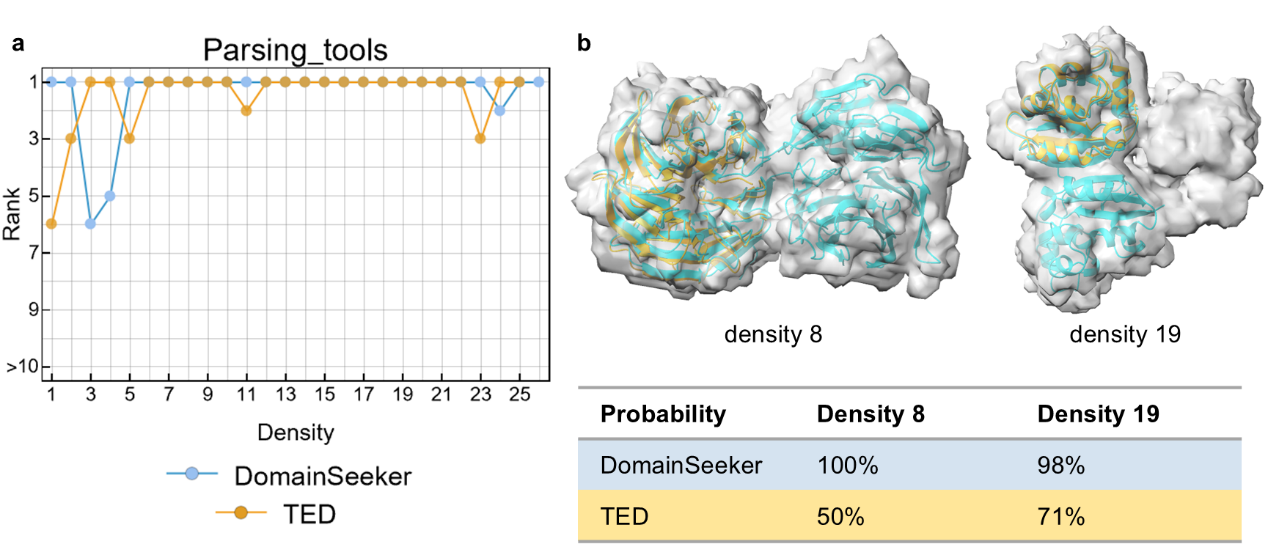

**Figure S7. Identification results using domains parsed by DomainSeeker versus TED**

DomainSeeker results are shown in cyan and TED results in orange. TED (the consensus of Chainsaw, Merizo, and UniDoc, now incorporated into AFDB) is designed to identify structurally or functionally annotated domains, whereas DomainSeeker defines domains as maximally large, internally rigid units optimized for density fitting. (a) Overall ranking performance on the sperm MIP dataset is similar between the two approaches. For density26, the correct domain given by TED falls below the minimum domain size threshold of 40 residues, so no valid result was produced. (b) Selected cases (densities 8 and 19) illustrating the characteristic difference: TED tends to produce smaller, function-oriented domains, while DomainSeeker merges regions into larger rigid units when PAE-based consistency permits. In both cases, the DomainSeeker-parsed domains yield substantially higher prior probabilities than their TED counterparts, reflecting stronger structural signals and improved discrimination between similar candidates.

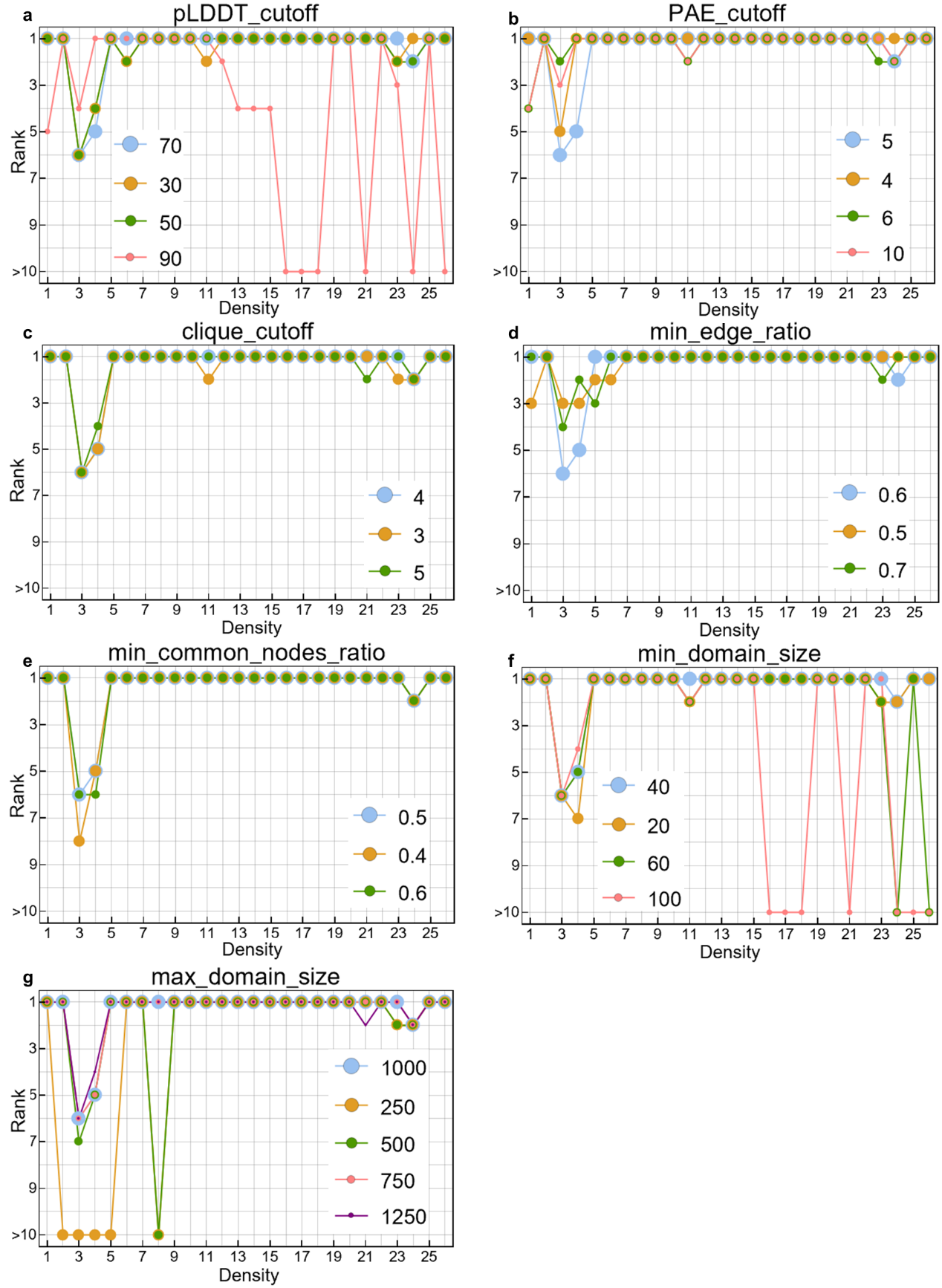

**Figure S8. Robustness to domain parsing parameters.**

Rankings of correct domains at varied parameters. (a) pLDDT_cutoff: 30, 50 and 70 give nearly identical results, while 90 drops accuracy substantially by losing structural continuity and identifiable features. (b) PAE_cutoff, (c) clique_cutoff, (d) min_edge_ratio, (e) min_common_nodes_ratio: moderate variations have limited impact on overall performance. (f) min_domain_size and (g) max_domain_size: overly restrictive settings (min ≥ 60 or max ≤ 500 residues) can exclude correct domains; results remain stable otherwise.

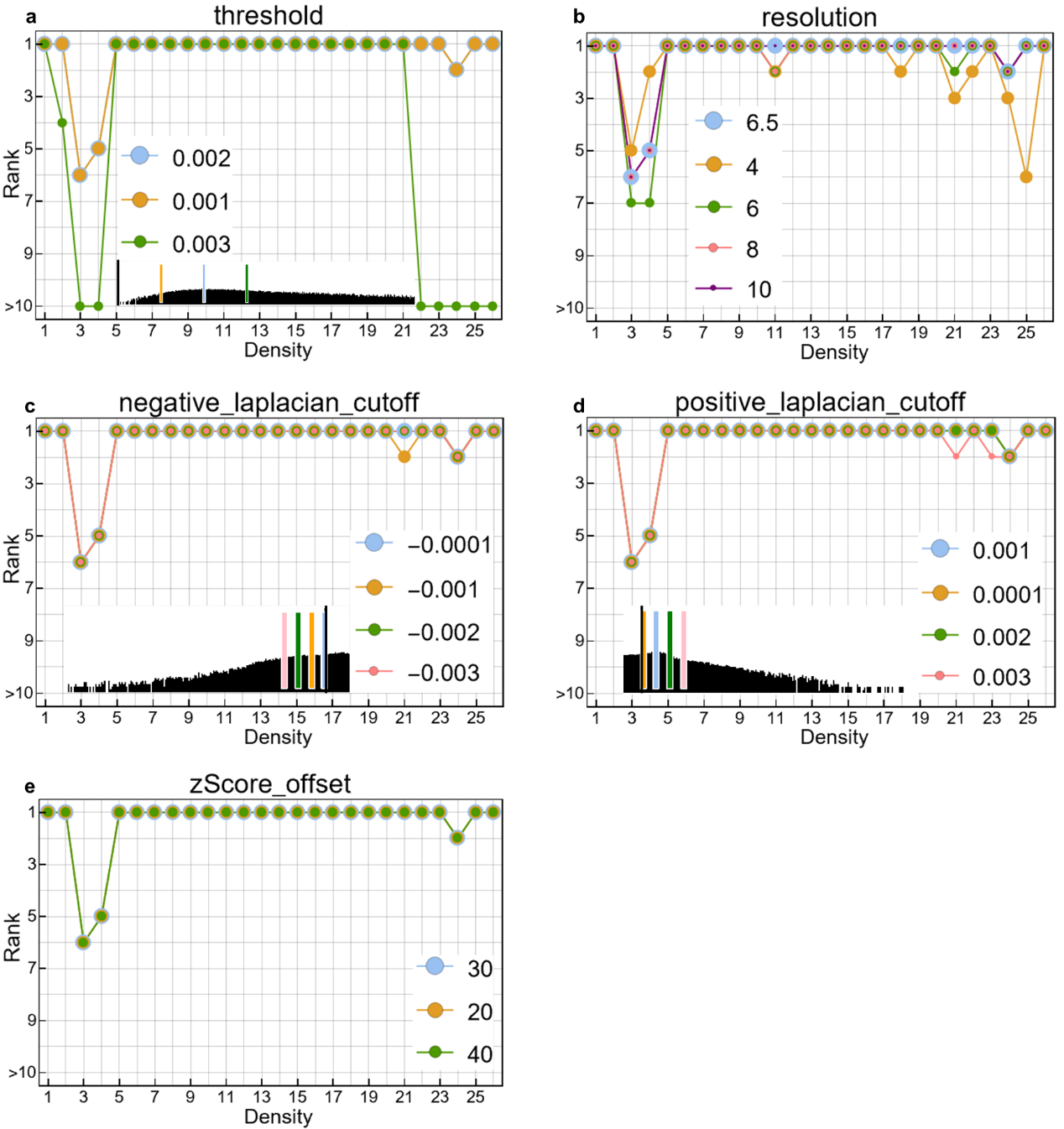

**Figure S9. Robustness to fitting and scoring parameters.**

Rankings of correct domains under varied fitting and scoring parameters on the sperm MIP dataset. (a) threshold: the density display and calculation threshold used for fitting and scoring. (b) resolution: the estimated resolution of the input map—identification accuracy is largely unaffected unless the input resolution deviates substantially from the true resolution. (c) negative_laplacian_cutoff and (d) positive_laplacian_cutoff: cutoffs used to suppress noise in the Laplacian-filtered map. Panels (a), (c) and (d) show the density histogram of density 1, with the corresponding level positions marked and the level 0 indicated by a black bar. In these three cases, identification remains stable as long as the parameters remove obvious noise while adequately covering the domain. (e) zScore_offset: prevents extreme dominance of the prior probability by excessively large z-scores, which may arise from insufficient data or other numerical artifacts. When z-scores are estimated accurately, this offset has negligible effect on the results as long as it exceeds the maximum z-score.

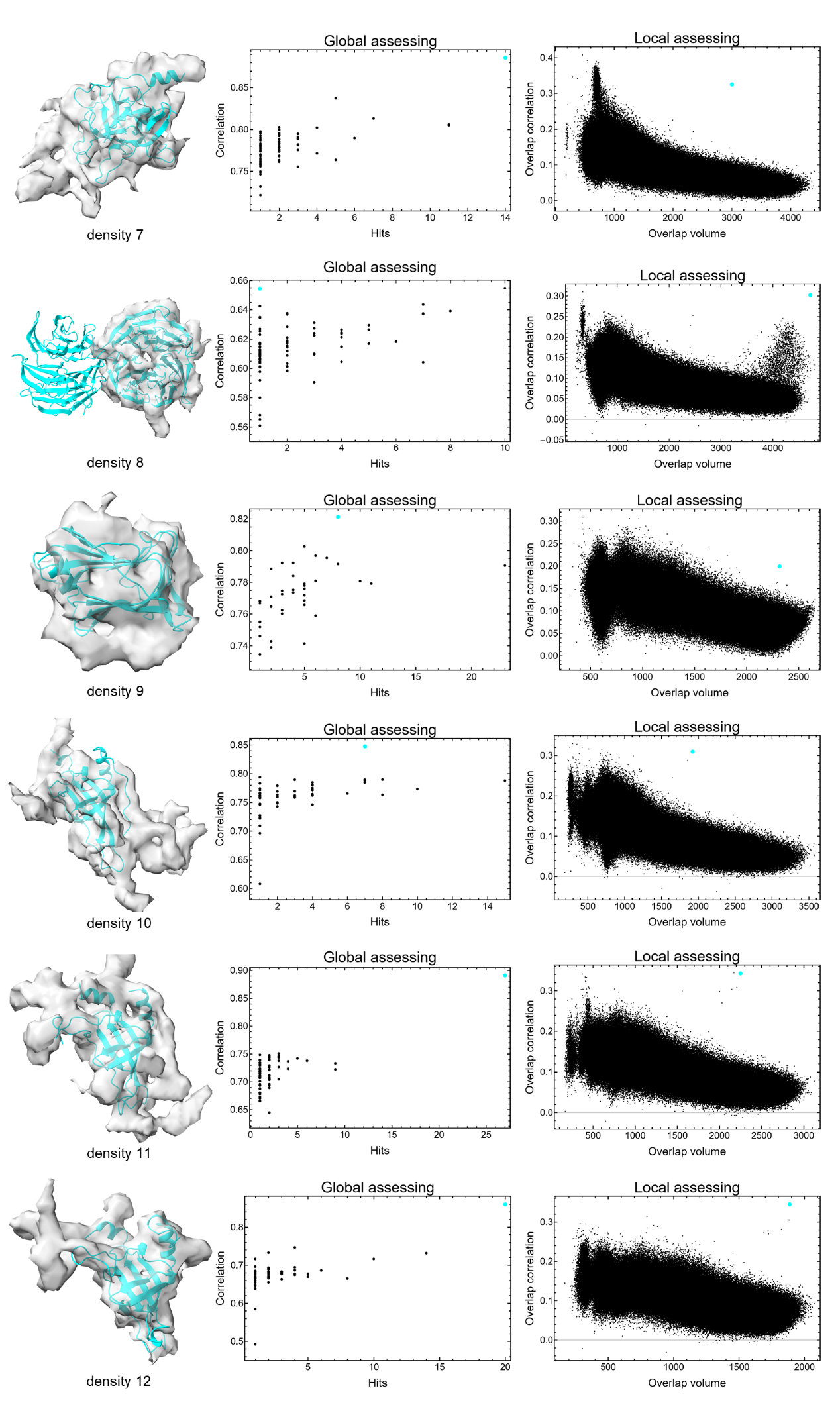
**(Continued on the next page)**

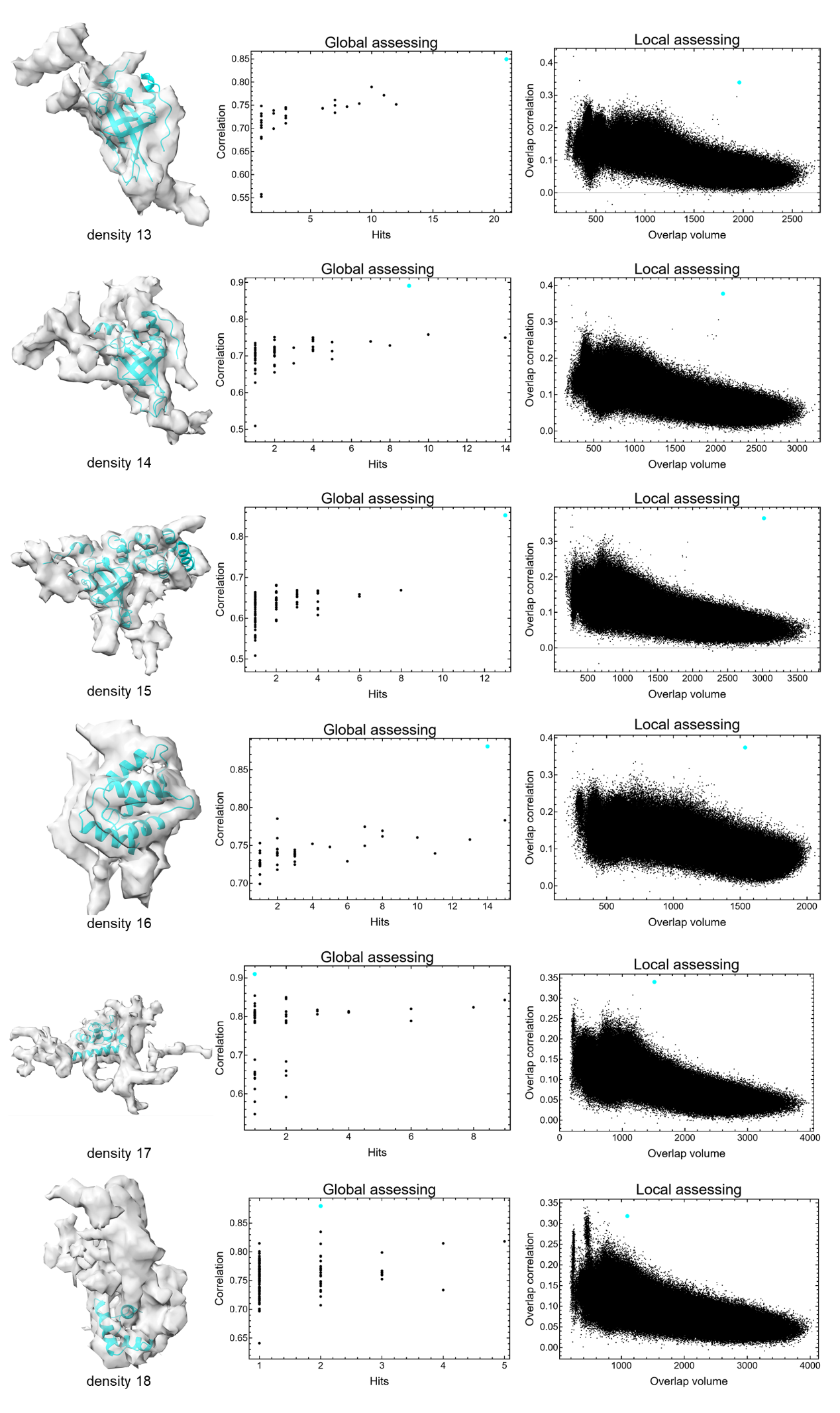
**(Continued on the next page)**

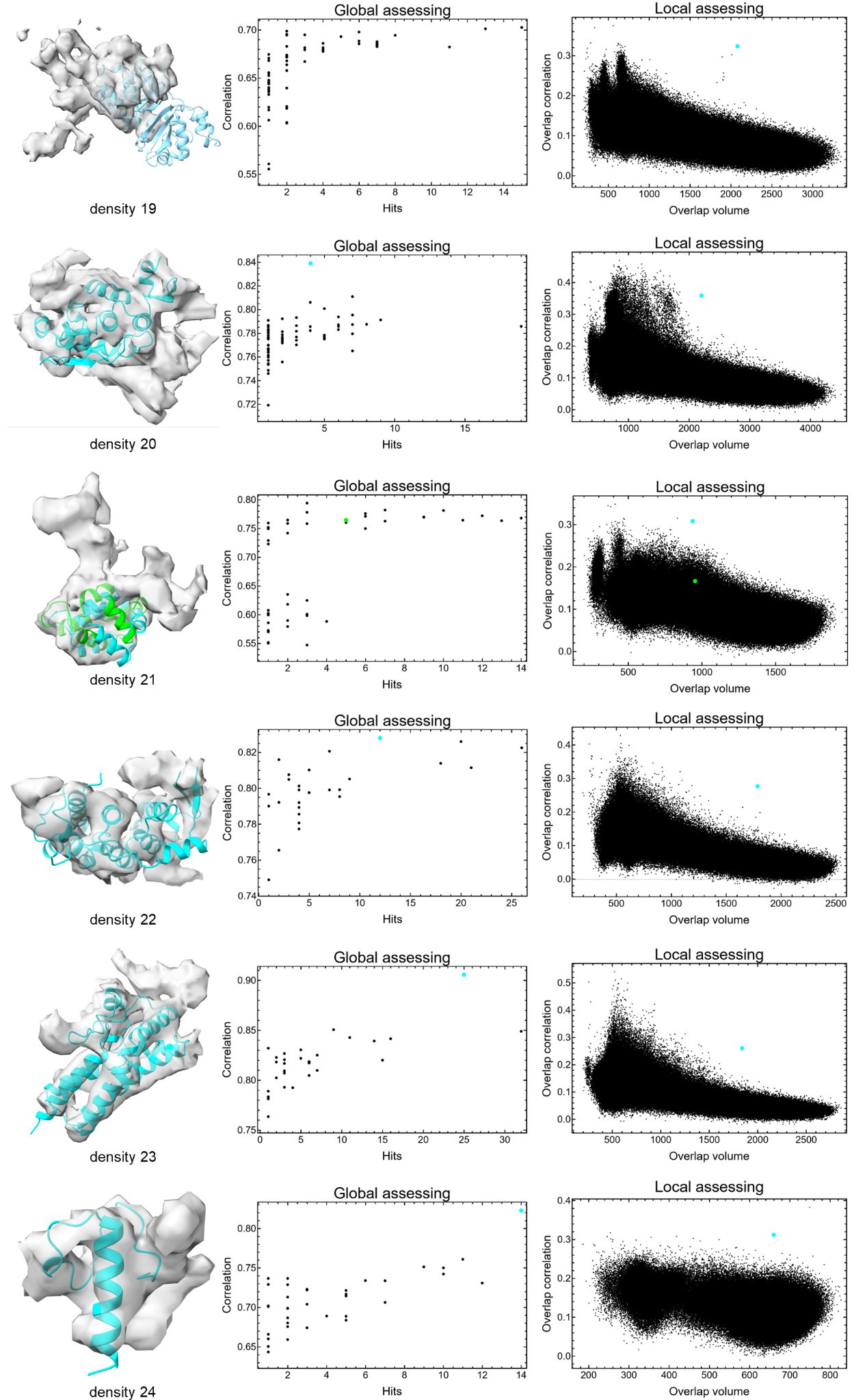

**(Continued on the next page)**

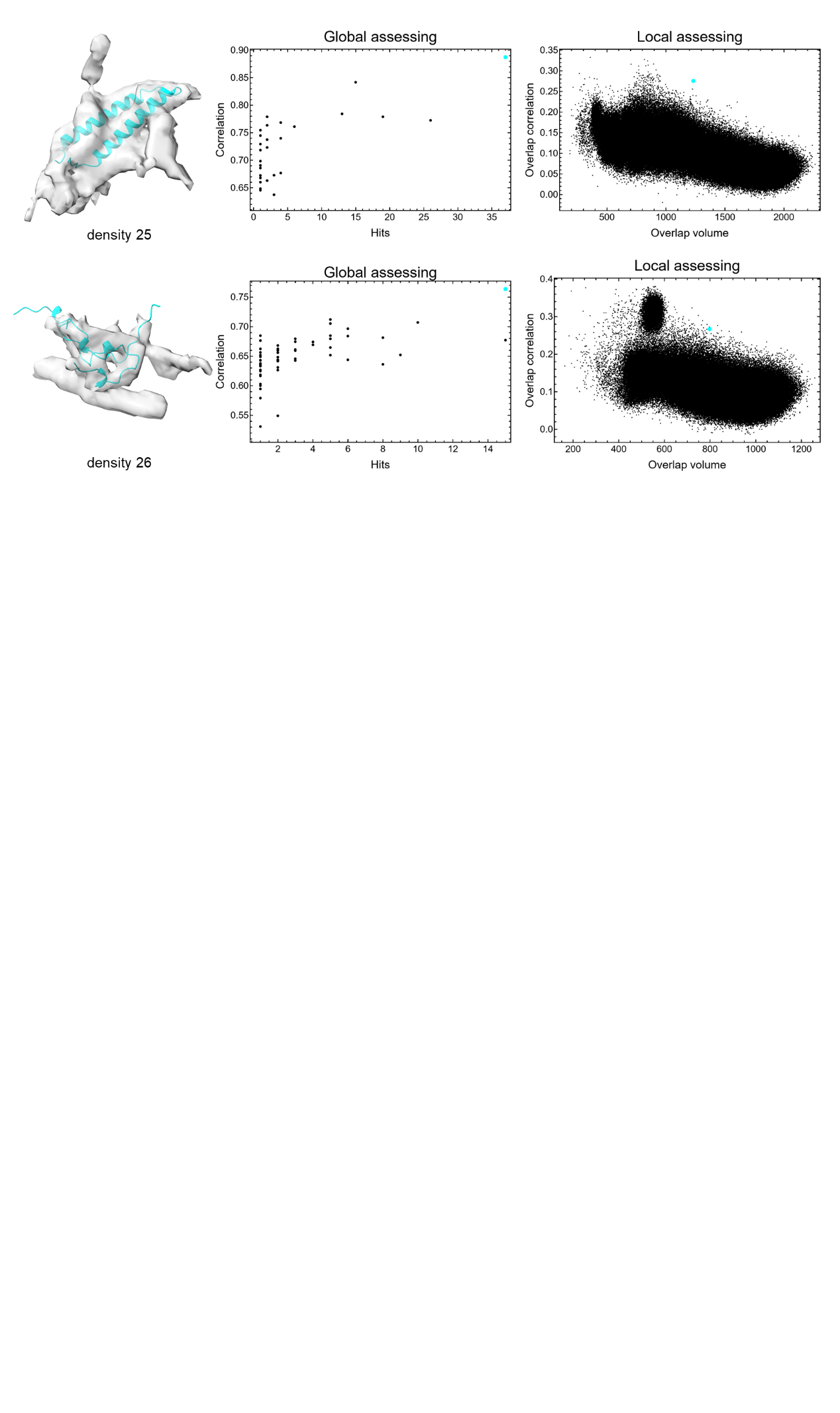

**Figure S10. Intermediate results of DomainSeeker on auto-segmented densities at 6.5** **Å from the mouse sperm microtubule doublet**

Each row represents a density. The left panel: the correct fitting position of the correct domain (cyan structure). The middle panel: hits and correlation values of candidate fittings for the correct domain (the cyan point indicates the correct position). The right panel: overlap volumes and correlations of all domain fittings (the cyan point marks the correct fit). For densities 19 and 21, the cyan structures/points indicate the correct fitting positions, manually added and absent from ChimeraX's candidates. The plots show that had these positions been included, DomainSeeker would have identified them correctly. For density 19, the segmented density excluded a large portion of the structural domain, so ChimeraX did not recognize the correct fit as a valid fitting. For density 21, the correct fit structure differed substantially from the structure at the local maximum identified by ChimeraX (green). For density 26, the domain is intrinsically small. Slight misalignment—whether redundancy or deficiency—in the segmented density relative to the domain boundary substantially alters its identifiable features, preventing the local z-score from standing out sufficiently for reliable identification.

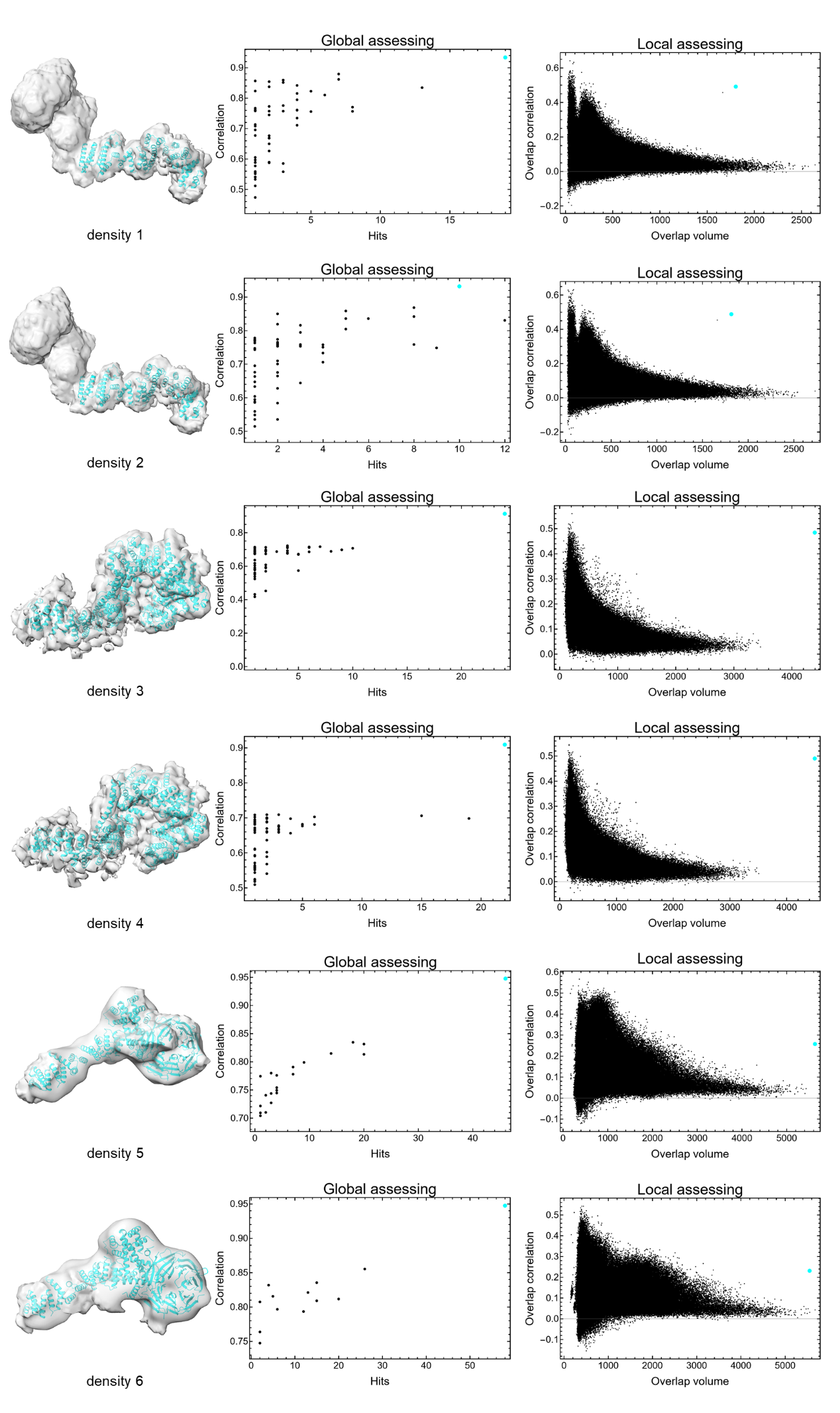
**(Continued on the next page)**

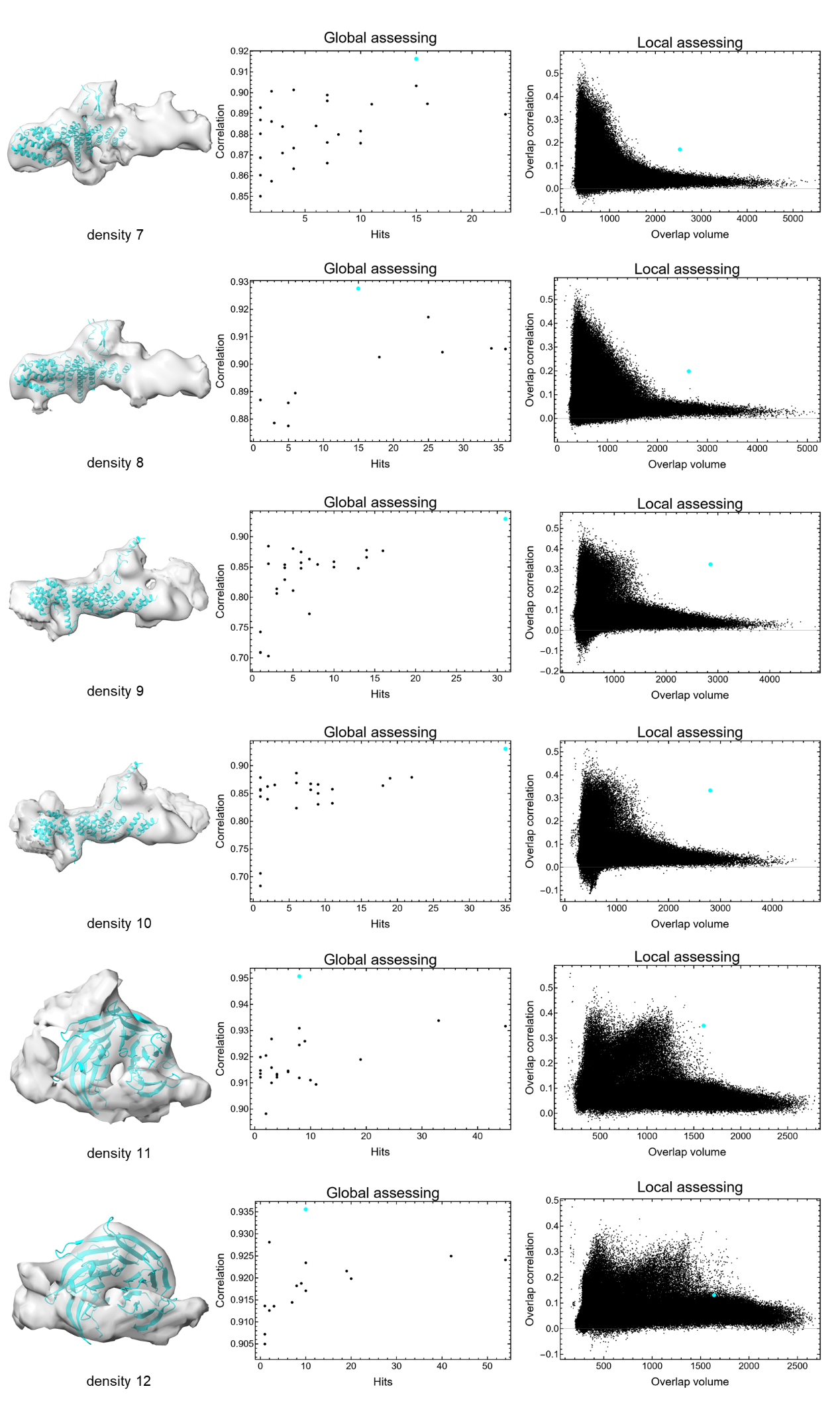
**(Continued on the next page)**

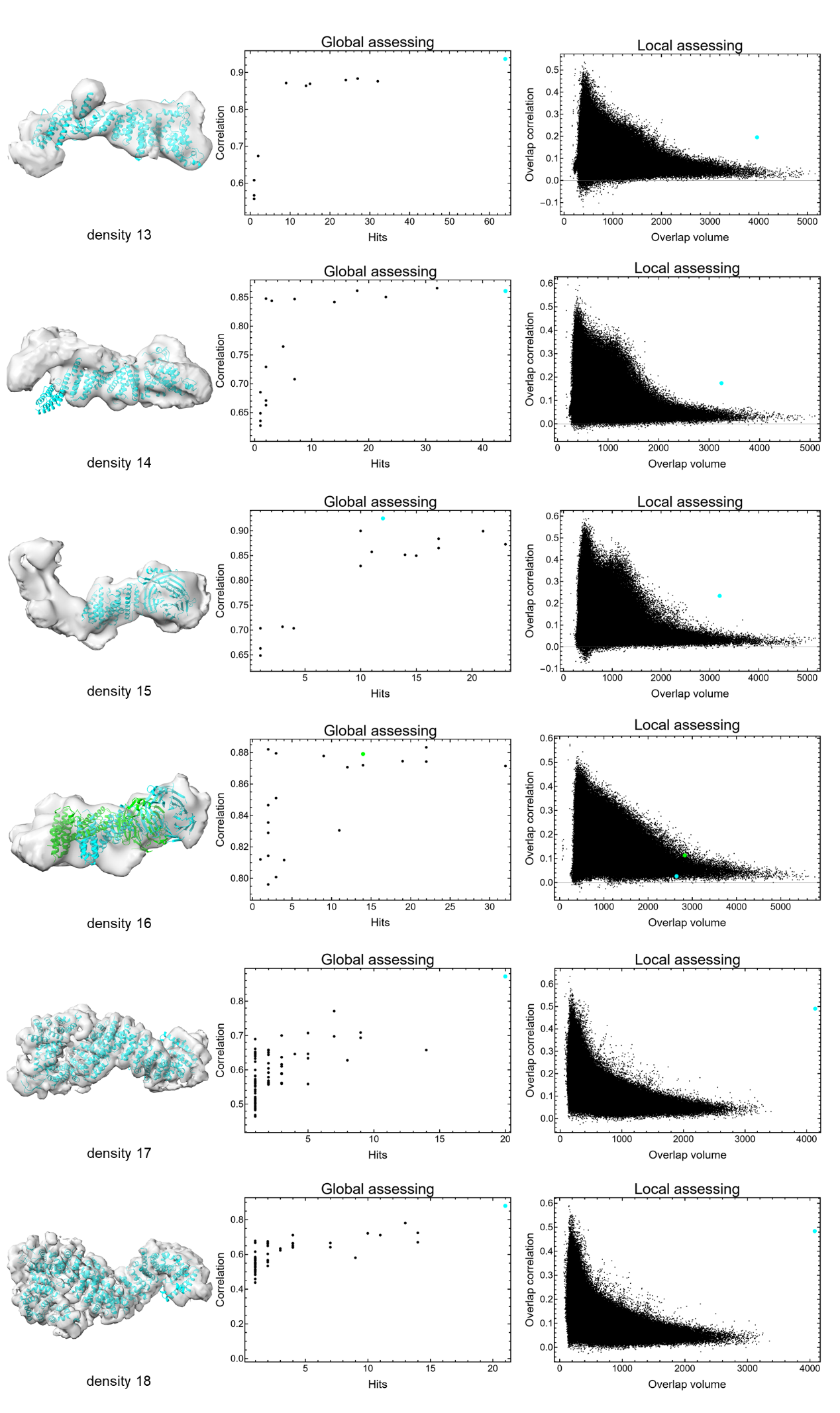
**(Continued on the next page)**

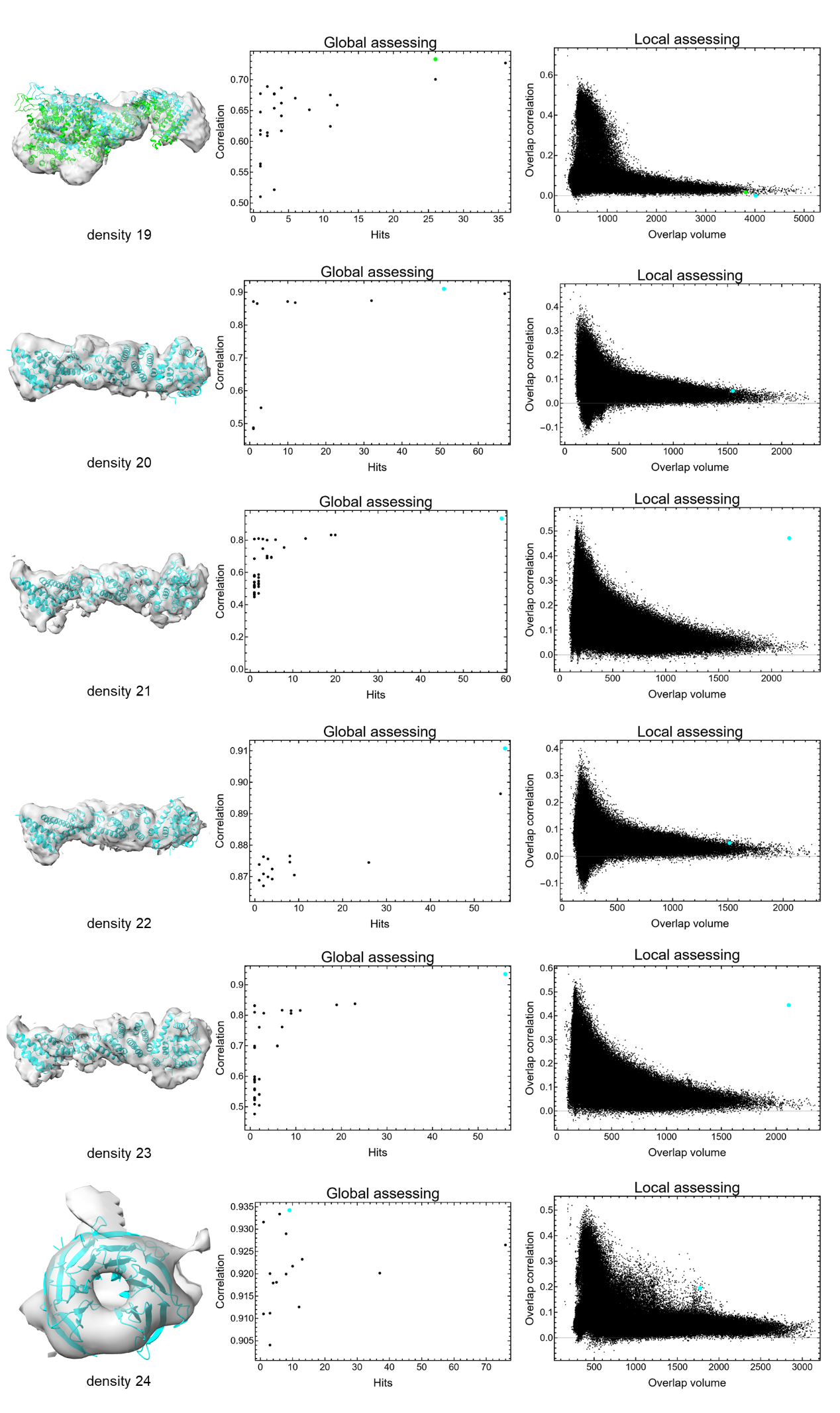
**(Continued on the next page)**

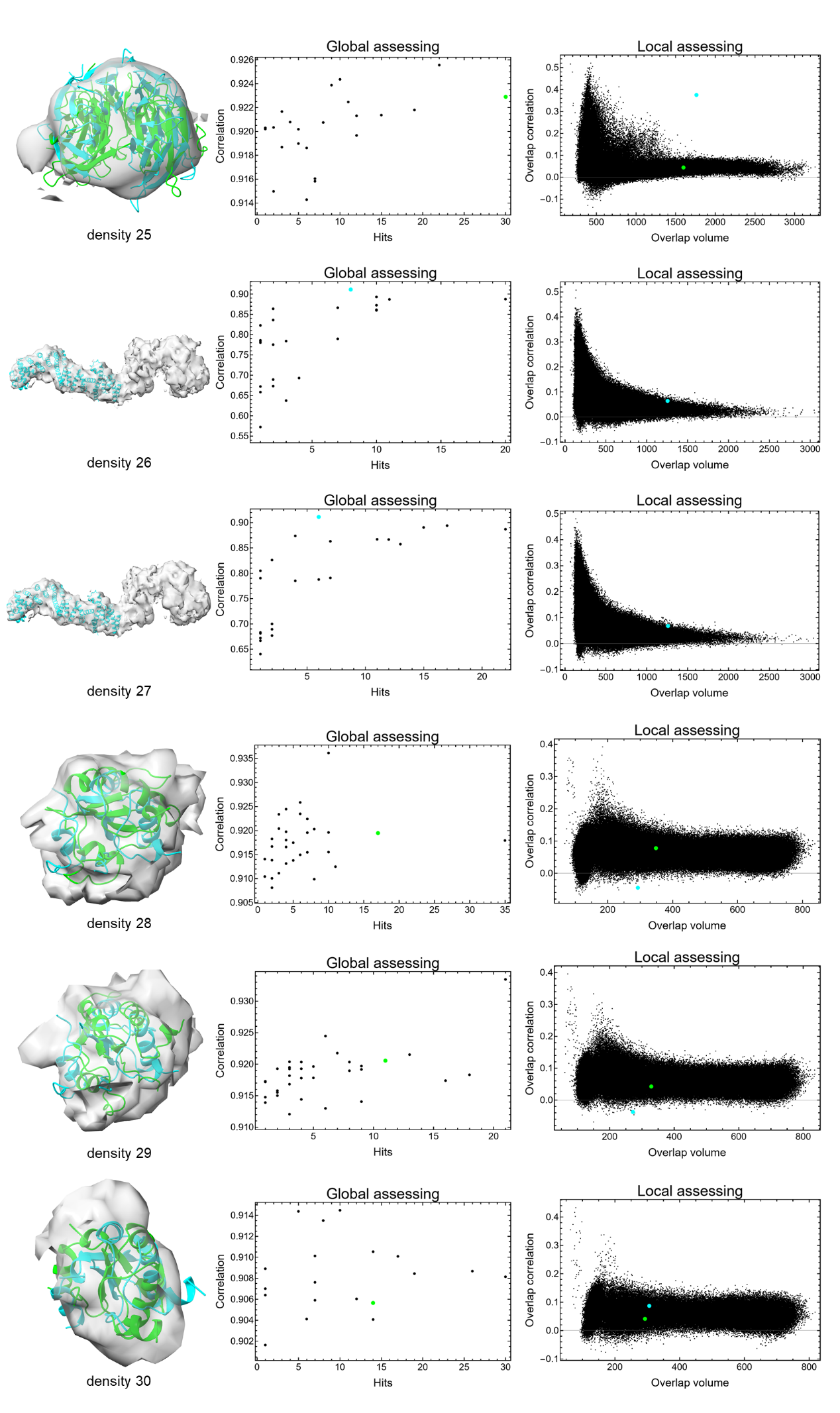
**(Continued on the next page)**

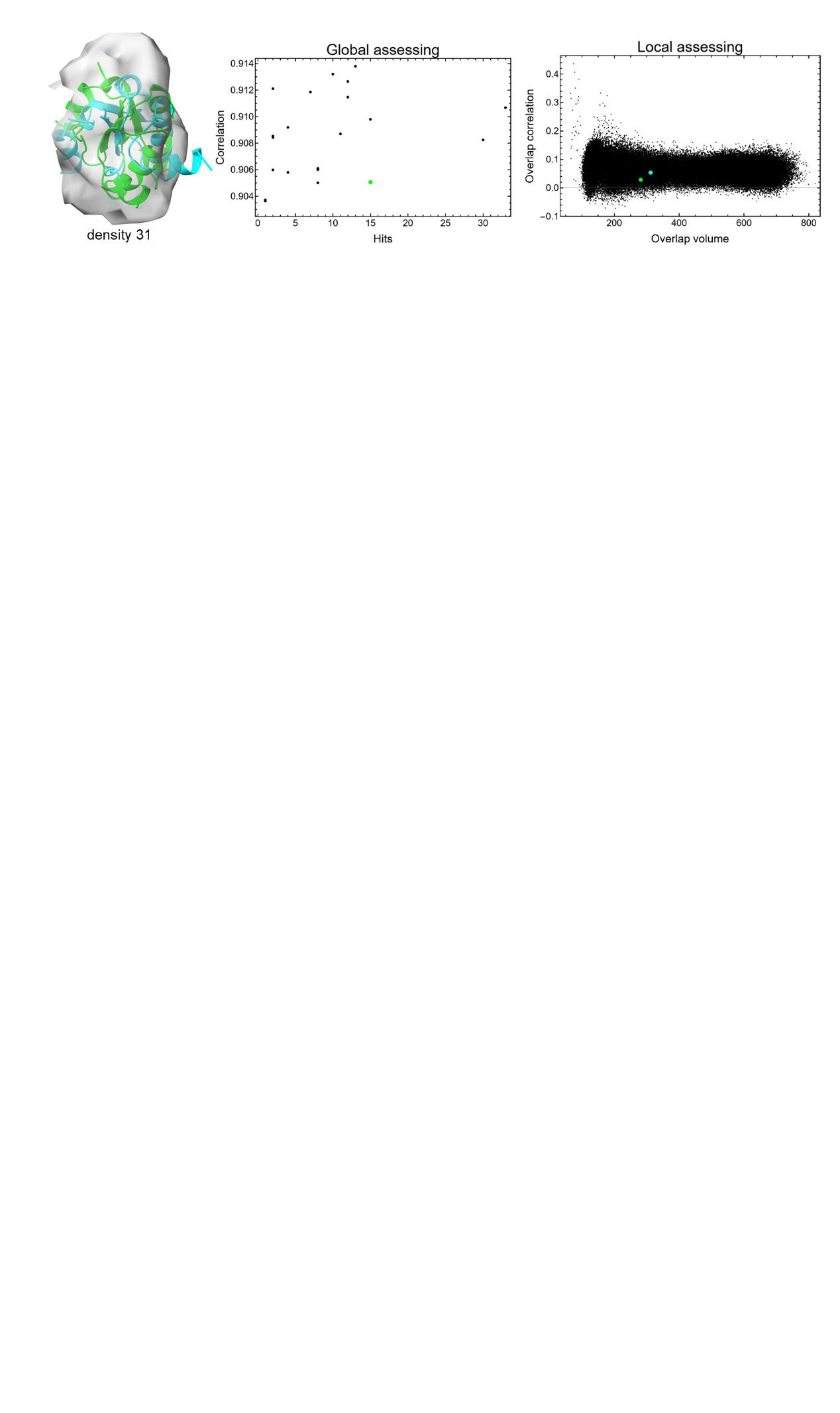

**Figure S11. Intermediate results on densities segmented from the yeast nuclear pore complex.**

Each row represents a density. The left panel: the correct fitting position of the correct domain (cyan structure). The middle panel: hits and correlation values of candidate fittings for the correct domain (the cyan point indicates the correct position). The right panel: overlap volumes and correlations of all domain fittings (the cyan point marks the correct fit). For densities 16, 19, 25 and 28-31, the cyan points (the correct fits, yet not the local correlation maxima) in local assessment plots are manually added (not in the raw data); The green points/structures denote the local maxima.

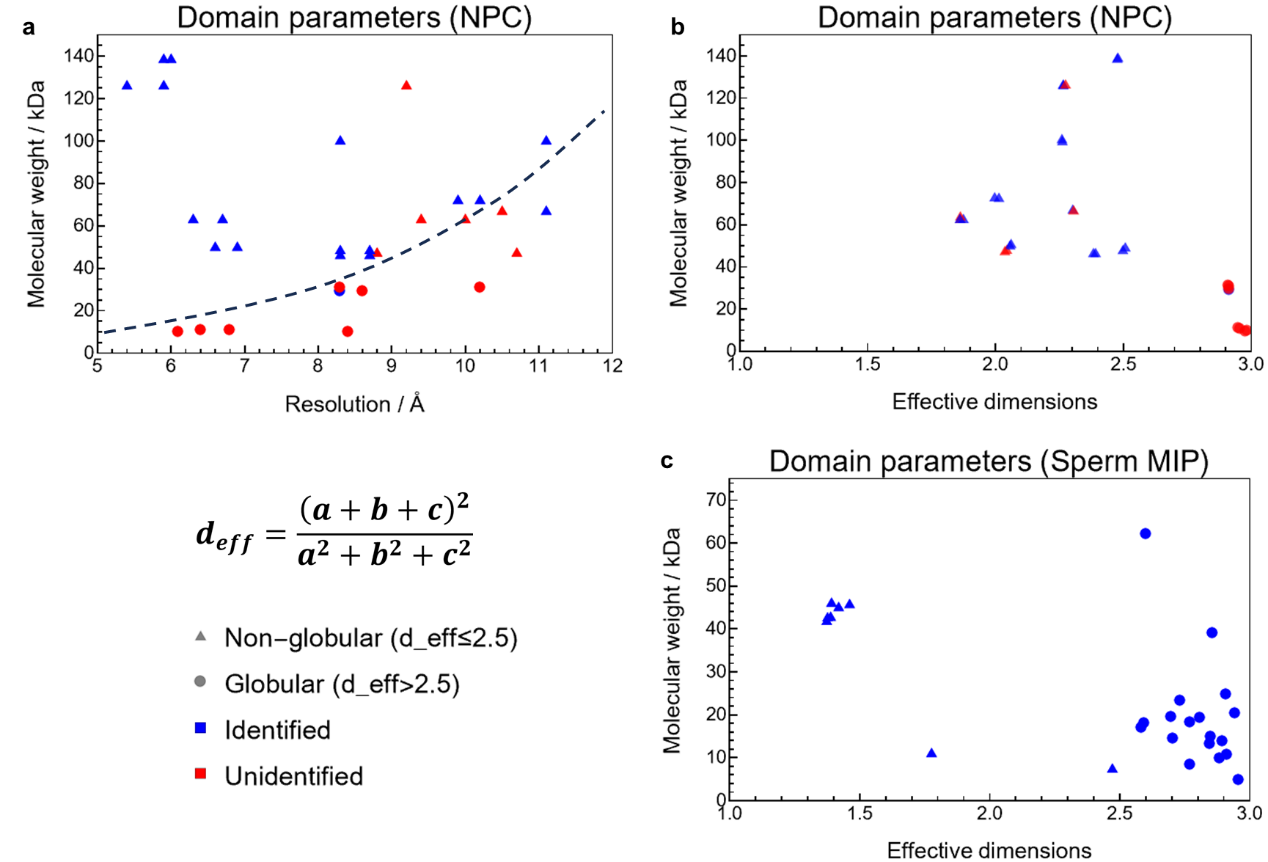

**Figure S12. Characterization of domain diversity across benchmark datasets.**

The effective dimensionality of each domain is quantified as $d_{eff}=\left( a+b+c \right)^{2}/(a^{2}+b^{2}+c^{2})$, where a, b, and c are the principal semi-axes of the inertia ellipsoid, with values ranging from 1 (rod-like) to 3 (globular). Circles mark globular domains (d_eff > 2.5) and triangles mark non-globular domains (d_eff ≤ 2.5). Blue and red indicate successfully identified and missed domains, respectively. (a-b) Yeast NPC dataset. Identification success is correlated with resolution, molecular weight and shape: correctly identified domains tend to have higher resolution, larger molecular weight and non-globular shape. The dashed line marks an approximate boundary between successful and failed identification. The overall trend of the boundary line suggests that poorer resolution requires larger molecular weight for domain identification. (c) Sperm MIP dataset (~6.5 Å). These identified globular domains demonstrate that the difficulty with compact domains observed in panel (a,b) is resolution-dependent rather than an intrinsic limitation of DomainSeeker. Overall, across both datasets, the tested domains span effective dimensionalities from 1 to 3, molecular weights from 4 to 150 kDa, and local resolutions from 5 to 11 Å, establishing that the method has been evaluated across diverse domain types varying in size, shape, and resolution.

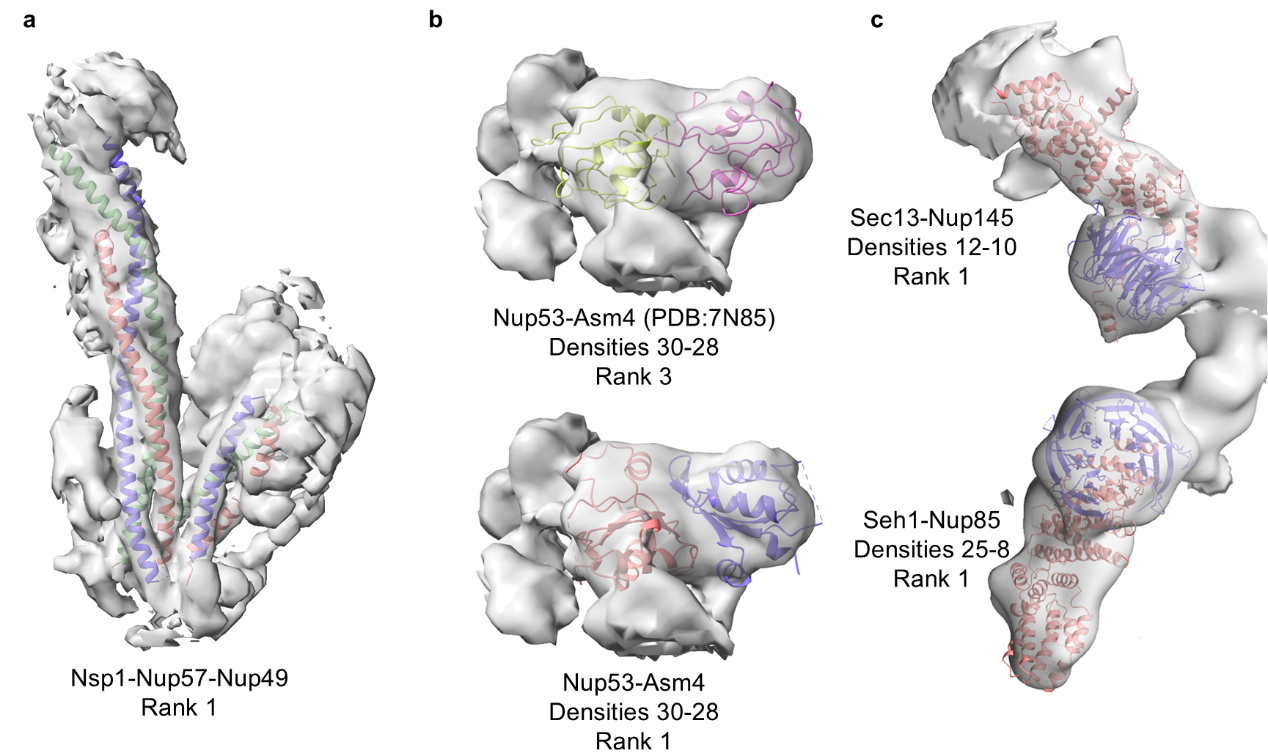

**Figure S13. Multimers identified by DomainSeeker.**

AF3-predicted structures and PAE data were used as input for DomainSeeker's domain parsing, and certain regions spanning multiple proteins were identified as single domains. DomainSeeker considers the relative positions and orientations among these regions to be reliable, and successfully identified some cross-protein domains from the densities corresponding to several complexes. In this figure, each density region is fitted with a domain composed of fragments from different proteins (rendered in distinct colors). (a) Nsp1–Nup57–Nup49. Individually, each protein consists of multiple single helices, making domain-level identification difficult. However, they form a complex with rigid domain-like segments, and DomainSeeker ranks the correct complex domain first. (b) Nup53–Asm4, a complex formed by two small globular domains. Neither domain could be identified alone (NPC densities 30 and 28). The predicted interface of the complex differs from the reference structure (PDB 7N85). The predicted structure ranks first, and the reference structure ranks third. (c) Sec13–Nup145 and Seh1–Nup85, each consists of one large and one small domain. The density corresponding to the smaller domain is at low resolution, precluding its independent identification (NPC densities 12 and 25). Once formed into cross-protein domains, these complexes are successfully identified by DomainSeeker as intact units.

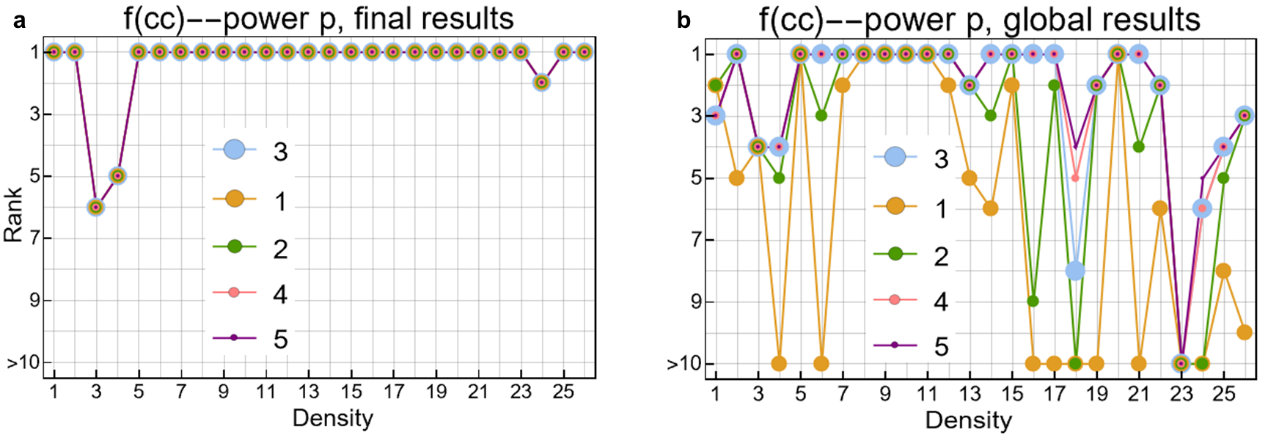

**Figure S14. Effect of the correlation transformation exponent on identification.**

(a) Effect of the exponent p in $f(cc) = \left( \frac{\mathrm{cc}}{1-cc} \right)^{p}$ under the combined scoring function. (b) Effect of p under the global scoring alone: higher exponents drive the global probabilities of high-correlation fits closer to 1, improving some rankings. Among them, p = 3 provides a trade-off between sensitivity and noise robustness.
